# Efficiency is better than speed for competitive suppression of drug-resistance

**DOI:** 10.64898/2026.05.29.728870

**Authors:** Clayton E. Cressler, Jessica L. Hite

## Abstract

Antimicrobial resistance (AMR) is a growing threat to human health, agriculture, and natural ecosystems. While antimicrobial use is a primary driver of resistance evolution, growing evidence suggests that anthropogenic environmental change may also contribute to the emergence and spread of resistant microbes. Yet the mechanisms linking environmental conditions to AMR remain poorly understood and are largely absent from current strategies to manage resistance. This gap arises, in part, because we still know surprisingly little about how ecological interactions shape the evolutionary dynamics of antimicrobial resistance. Here, we develop a general eco-evolutionary modeling framework to investigate how spatial structure, resource availability, and growth-efficiency trade-offs shape competition between drug-sensitive and drug-resistant strains. We show that spatial heterogeneity influences resistance evolution not simply by strengthening or weakening competition, but by altering the ecological mechanisms through which competition suppresses resistant strains. In homogeneous environments, resistance is suppressed primarily by faster-growing drug-sensitive competitors and only under a relatively narrow range of ecological conditions. In contrast, heterogeneous environments with limited resources favor more resource-efficient drug-sensitive strains, which outcompete resistant strains through a distinct colonization advantage that operates across a broader region of parameter space. These patterns emerge regardless of whether resistant mutants are initially present or arise *de novo* through mutation. Together, our results provide insight into mechanistic links between environmental change and AMR emergence, suggesting that environmental homogenization and resource enrichment may reduce opportunities for competitive suppression and thereby increase resistance risk. More broadly, this work demonstrates how ecological interactions and environmental structure shape the evolution of resistance and offers possible insight into resistance management.

**Author summary:** Antimicrobial resistance (AMR) is often viewed as a direct consequence of antimicrobial use. However, mounting evidence suggests that ecological processes may often suppress resistant strains before they can emerge and spread. Despite growing recognition that ecological interactions influence AMR, we still know relatively little about the environmental conditions that strengthen or weaken those interactions. We developed a general eco-evolutionary modeling framework that integrates consumer-resource theory, spatial ecology, and stochastic trait evolution to investigate how spatial structure, resource availability, and growth-efficiency trade-offs shape competition between drug-sensitive and drug-resistant strains. Whereas previous theoretical and empirical work has largely focused on spatial variation in antimicrobial exposure, our framework examines how spatial variation in resource availability interacts with growth-efficiency trade-offs to influence the evolutionary dynamics of AMR. Our analyses reveal that environmental heterogeneity does not simply strengthen or weaken competition; it fundamentally alters the ecological mechanisms through which competition suppresses resistance. In homogeneous environments, resistance is constrained primarily by faster-growing sensitive strains but only under a relatively narrow range of ecological conditions. In contrast, heterogeneous environments favor more resource-efficient competitors that suppress resistant strains through a distinct colonization-based mechanism that operates across a larger region of parameter space. The key takeaway is not just that spatial structure or resource availability matter for AMR. Instead, the main lesson is that environmental homogenization may help promote AMR. Because many human activities homogenize environments (e.g., invasive plants, chemically intensive agriculture typical across much of the US Corn Belt), it is critical to determine whether they do so at the spatial scales relevant to microbial competition. Future studies integrating AMR surveillance data and environmental metadata with this eco-evolutionary model could help move environmental heterogeneity and microbial community structure from correlates of resistance into mechanistic predictors of AMR.

## Introduction

Why do some hosts and environments emerge as ‘hotspots’ of antimicrobial resistance (AMR), while others do not? Until recently, efforts to address this question have focused primarily on antimicrobial exposure as the dominant driver of resistance evolution. However, we now realize that while antimicrobial exposure strongly selects for resistance, this metric alone cannot explain the striking variation in resistance observed in livestock systems or environmental reservoirs [1–4]. The ubiquity of this pattern underscores the need to identify other key factors that shape the evolution of antimicrobial resistance [5–7].

At least part of the answer lies in the ecological processes that determine the competitive success of resistant pathogens. Although antimicrobial resistance is a trait of individual bacteria, whether it evolves in populations exposed to antimicrobials is determined, in part, by the complex microbial communities in which pathogens live, grow, and compete [8–10]. In the absence of antimicrobial exposure, resistant strains are often competitively suppressed by drug-sensitive strains, particularly when resistance incurs a fitness cost [11, 12]. Antimicrobial treatment can relax this suppression by reducing susceptible competitors, allowing resistant pathogens to proliferate through competitive release [13–15]. This suggests competition, and perhaps other ecological interactions, could be strategically leveraged to help manage antimicrobial resistance [11, 16, 17]. Indeed, mounting empirical and theoretical evidence suggests that ecological interactions can shift the trajectory of resistance evolution [18–21].

Despite these exciting findings, disease management based on such assumptions could quickly go astray without accounting for other factors like resource availability that govern ecological processes like competition, distribution, and diversity [15, 22]. For instance, drug-resistant and drug-sensitive strains can compete for limiting resources such as target cells, iron, and glucose [23–26]. Theory predicts that when organisms compete for a single limiting resource, the organism able to persist at lower resource levels will outcompete its competitor [27–30]. Hence, when resistance is associated with elevated resource requirements, limiting resource availability can give drug-sensitive strains a competitive advantage, effectively preventing the emergence of resistance [16, 23, 31, 32]. In contrast, resource enrichment and high-dose drug therapy can jointly prevent competitive suppression and fuel resistance. An elegant experiment in malaria parasites demonstrates this possibility: increasing resource availability enabled resistant parasites to rapidly overcome competitive suppression following drug treatment and, in some cases, achieve higher densities in the presence of competitors than in their absence [33]. Resource availability could, therefore, prove especially important to resistance management because it may enhance or diminish competitive suppression.

Further complicating matters, competitive outcomes hinge not only on resource availability but also whether the distribution of those resources is homogeneous or heterogeneous, due to, for example, spatial structure. Spatial structure has long been recognized as an important determinant of evolutionary dynamics [**?**, 34–36]. Across ecology and evolutionary biology, theoretical and empirical studies demonstrate that spatial structure can alter rates of adaptation, maintain diversity, and promote coexistence by generating heterogeneity in abiotic and biotic factors that can shift interactions among competing genotypes [**?**, 28, 37, 38]. Similar ideas have been applied to antimicrobial resistance, although the predicted effects are notably mixed; see [39] and references therein. Some studies suggest that spatial structure can slow or inhibit the evolution of resistance [19, 40]. In other cases, spatial heterogeneity can accelerate adaptation when antimicrobial exposure varies across space [41–43] or when biofilms provide spatial refuges that facilitate the accumulation and persistence of resistant mutants [44]. These contrasting findings likely reflect differences in both the ecological assumptions and mechanisms incorporated into existing experimental and theoretical models.

In particular, most studies have focused on competition between a resistant mutant and its ancestral drug-sensitive strain, implicitly assuming that competitors differ primarily in their resistance phenotype [11, 12]. However, drug resistance evolves in organisms that vary across a broader suite of traits affecting competitive success. growth rate and resource-use efficiency (yield) because they are fundamental components of microbial fitness widely linked with adaptation and evolution [45–47]. The importance of growth-efficiency trade-offs in modulating competitive outcomes has a long history in ecological theory [27–30]. Despite this rich ecological foundation, how such trait variation shapes resistance evolution remains poorly understood [48–50].

Our goal here is to help advance this endeavor by developing a general eco-evolutionary modeling framework that integrates consumer-resource theory, spatial ecology [27–30], and stochastic trait evolution [51, 52]. Whereas previous theoretical and empirical work has largely focused on spatial variation in antimicrobial exposure, our framework examines how spatial variation in resource availability interacts with growth-efficiency trade-offs to influence competitive suppression of AMR.

We build understanding of our main finding - that more efficient strains are much better able to suppress the emergence of antimicrobial resistance - through a series of theoretical and computational explorations of increasing complexity. We begin by examining competition in a spatially homogeneous environment to develop intuition for how growth rate and growth efficiency affect resource and microbial equilibria. We then consider ecological competition in a spatially heterogeneous environment between drug-resistant and drug-sensitive strains with fixed differences in growth rate, growth efficiency, and drug-induced mortality rate to develop intuition about the environmental and competitive circumstances where drug-resistance is most likely to emerge. Finally, we test that intuition by integrating evolution into our ecological model and allowing drug-resistance to evolve dynamically.

To do so, we extend our previous work developing Gillespie eco-evolutionary models (GEMs) [51, 52]. GEMs use the Gillespie algorithm to create stochastic simulations of an underlying ecological model. Then, GEMs incorporate trait variation, mutation, and inheritance, allowing eco-evolutionary dynamics to emerge from simulating the stochastic processes of birth, death, and mutation [53].

## Methods Part 1: Model structure and competition scenarios

### Base model overview

We study competition between drug-sensitive (*S*_*s*_) and drug-resistant (*S*_*r*_) strains using a consumer-resource framework. We begin with a spatially homogeneous environment where the biomass dynamics of the two strains competing for a single limiting resource (*R*) are described by the following model:

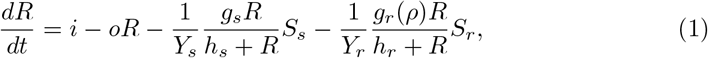

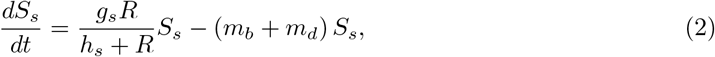

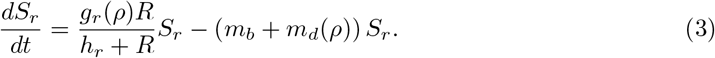

The resource (*R*) is assumed to follow semi-chemostat dynamics, with a constant resource influx at rate *i* and removal at rate *o*. Both strains uptake resource assuming Michaelis-Menten (Monod) uptake kinetics, with maximum growth rates *g*_*s*_ and *g*_*r*_ and half-saturation constants *h*_*s*_ and *h*_*r*_. The yield parameters *Y*_*s*_ and *Y*_*r*_ quantify the bacterial biomass produced per unit of resource biomass; although *Y* is typically defined as yield, it can be more intuitive to think of it as quantifying growth efficiency, with higher values of *Y* implying more efficient biomass production [47]. Bacterial biomass is lost at a background rate *m*_*b*_ and a drug-induced mortality rate *m*_*d*_.

### Growth–drug-resistance trade-off

It is commonly assumed that antimicrobial resistance is costly, often resulting in a reduction in growth rate [1, 12, 46]. We represent this cost by assuming that drug-resistance reduces drug-induced mortality at the expense of reduced growth in the resistant strain, *S*_*r*_. Specifically, we assume that this strain is characterized by an underlying resistance trait *ρ* that affects both its growth rate, *g*_*r*_(*ρ*), and its drug-induced mortality rate, *m*_*d*_(*ρ*). We assume the following functional relationships:

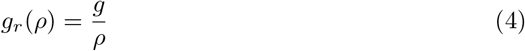

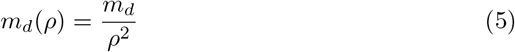

These functional forms capture a growth–resistance trade-off in which increasing resistance (higher *ρ*) reduces growth while lowering susceptibility to antimicrobials (drug-induced mortality). These functional forms were chosen because, in a spatially homogeneous environment, this trade-off gives rise to a single evolutionarily stable resistance strategy, 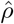 (see Supplementary Information for details of this derivation):

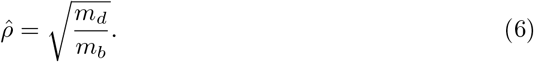

We use this evolutionarily stable trait value as a reference when parameterizing resistant strains in subsequent analyses.

### Competition scenarios

To evaluate where, when, and how competition can suppress the evolution of drug-resistance, we consider two competition scenarios. In the first scenario, we test whether a faster-growing but drug-sensitive strain can competitively suppress a resistant strain with a growth–resistance trade-off. This represents the more traditional view of competitive suppression, in which resistance carries a growth cost that leaves resistant strains vulnerable to competition from their faster-growing ancestors [11, 12]. Here, we assume that *g*_*s*_ *> g*_*r*_(*ρ*) for all values of resistance, *ρ*, and *m*_*d*_(*ρ*) *< m*_*d*_ according to the functional form defined above. This scenario allows us to assess whether a growth advantage alone is sufficient to suppress a drug-resistant competitor.

In the second scenario, we test whether competition from a resource-efficient drug-sensitive strain suppresses resistance more effectively than competition from a faster-growing strain. Although trade-offs between growth rate and resource-use efficiency are widespread in bacteria [47], their implications for competitive suppression of antimicrobial resistance remain largely unexplored. Here, we assume that *g*_*r*_(*ρ*) *> g*_*s*_ (at least for lower values of resistance, *ρ*) and that *Y*_*r*_ *< Y*_*s*_, so that the sensitive strain is more efficient at converting resources into bacterial biomass. As before, we assume that the resistant strain has *m*_*d*_(*ρ*) *< m*_*d*_ according to the functional form defined above. This scenario allows us to assess whether an efficiency advantage is sufficient to suppress a drug-resistant competitor.

We first consider a spatially homogeneous, single-patch environment to establish baseline expectations for both scenarios. We then layer in environmental heterogeneity and explore how that alters competitive outcomes. Finally, we examine how both environmental heterogeneity and resource availability interact to shape the evolution of drug-resistance.

## Methods Part 2: Single-patch outcomes

Before simulating competition in spatially structured environments, we first examine how variation in the growth advantage of a drug-sensitive strain (*γ*), the resource efficiency advantage of a drug-sensitive strain (*ξ*), and the drug-induced mortality rate (*m*_*d*_) shape baseline expectations for competition in a single, spatially homogeneous patch. More specifically, we analyze how changes in these parameters alter the equilibrium resource requirement 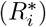 and equilibrium population density 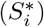attained by each strain when growing in a single-strain patch.

In consumer-resource theory, 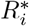 ratios predict competitive outcomes in homogeneous environments, with the strain characterized by the lower 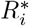 ratios expected to outcompete the other competitor [27, 29, 30]. This represents a tragedy of the commons: competition favors strains that most strongly exploit the shared resource. On the other hand, 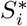 ratios determine which strain has a numerical advantage; while this does not matter for competition in a homogeneous environment, in a spatially structured environment the strain with a higher abundance may be more likely to colonize new environments, allowing it to persist and even dominate in competition [28, 38, 54]. Thus we predict that the strain with the lower 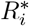 will win in a more homogeneous environment whereas the strain with the higher 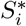 will win in a more heterogenous environment. We test each of these hypotheses with the assumption that the drug-resistant strain is at the ESS — evolutionarily stable strategy for drug-resistance,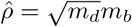.

### 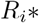 for drug-sensitive and drug-resistant strains

For the first scenario, we assume that the sensitive strain has a growth advantage given by *g*_*s*_ = *γg*, where *g*_*r*_ = *g* and *γ ≥* 1; and that the yield coefficients and half-saturation constants are the same for both strains (*Y*_*s*_ = *Y*_*r*_ = *Y, h*_*s*_ = *h*_*r*_ = *h*).

Here, the equilibrium resource levels for each strain are:

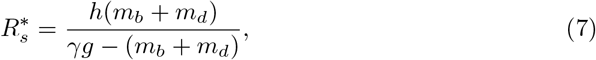

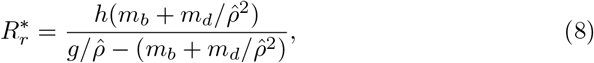

The drug-sensitive strain is predicted to outcompete the drug-resistant strain whenever 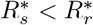 Assuming that the drug-resistant strain is at the ESS for drug resistance, 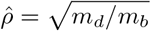 this condition simplifies to

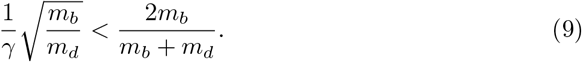

In this case, the competitive outcome depends on both the growth advantage of the drug-sensitive strain *γ* and the drug-induced mortality rate *m*_*d*_, with larger values of *γ* favoring the drug-sensitive strain.

For the second scenario, we assume that the sensitive strain has an efficiency advantage given by *Y*_*s*_ = *ξY*, where *Y*_*r*_ = *Y* and *ξ ≥* 1, that the resistant strain has a maximum growth rate that is ten times faster than the sensitive strain, so *g*_*r*_(*ρ*) = 10*g/ρ* and *g*_*s*_ = *g*, and that the half-saturation constants are equal (*h*_*s*_ = *h*_*r*_ = *h*).

Here, the equilibrium resource levels for each strain are:

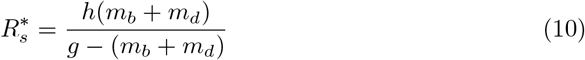

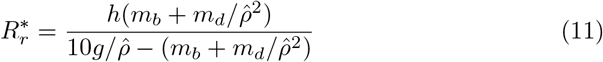

Because resource efficiency (*Y*_*i*_) does not appear in the expression for 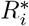, being more efficient provides no advantage in direct resource competition in spatially homogeneous environments. Hence, in this scenario, the outcome of competition depends only on the drug-induced mortality rate, *m*_*d*_. For the baseline parameters we use here *m*_*b*_ = 0.01, the efficient, drug-sensitive strain can never outcompete the drug-resistant strain.

### 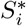for drug-sensitive and drug-resistant strains

For the first scenario, the equilibrium densities of each strain in a patch are

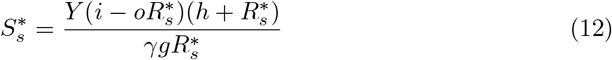

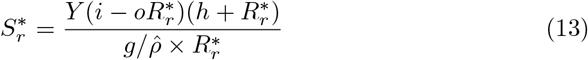

For the second scenario, the equilibrium strain densities are

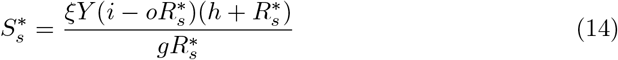

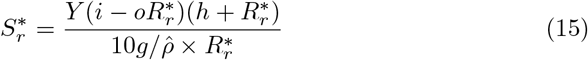

Although it is hard to gain much analytical insight here, given that strain equilibria 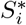 depend on resource equilibria 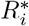,numerical calculation of these strain densities for a range of drug-induced mortality rates and either sensitive growth rates (scenario 1) or sensitive growth efficiencies (scenario 2) can give some insight into which strain is likely to have an advantage in a spatially heterogeneous environment. Thus, consideration of resource and strain equilibria in a single-patch, homogeneous environment, can help guide predictions for more complex, spatially heterogeneous environments.

## Results Part 1: Single-patch outcomes

Fig. 1 shows how changing the growth rate (*γ*) or growth efficiency (*ξ*) advantage of the drug-sensitive strain and the drug-induced mortality coefficient (*µ*) affects the single-strain equilibria for the drug-sensitive and drug-resistant strains. That is, for each value of growth advantage and mortality coefficient (panels A and B) and for each value of efficiency advantage and mortality coefficient (panels C and D), we calculate the ratio of the single-patch, single-strain resource equilibria 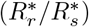(panels A and C) and the strain abundance equilibria 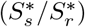 (panels B and D). Note that these equilibria are calculated assuming each strain is present on its own, rather than in competition. Then, their ratio is taken to facilitate predictions about which strain has a competitive advantage (lower 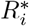) and which has a colonization advantage (higher 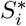). Note also that which strain is in the numerator changes in the two ratios so that larger numbers will always indicate an advantage for the drug-sensitive strain.

**Fig 1.**
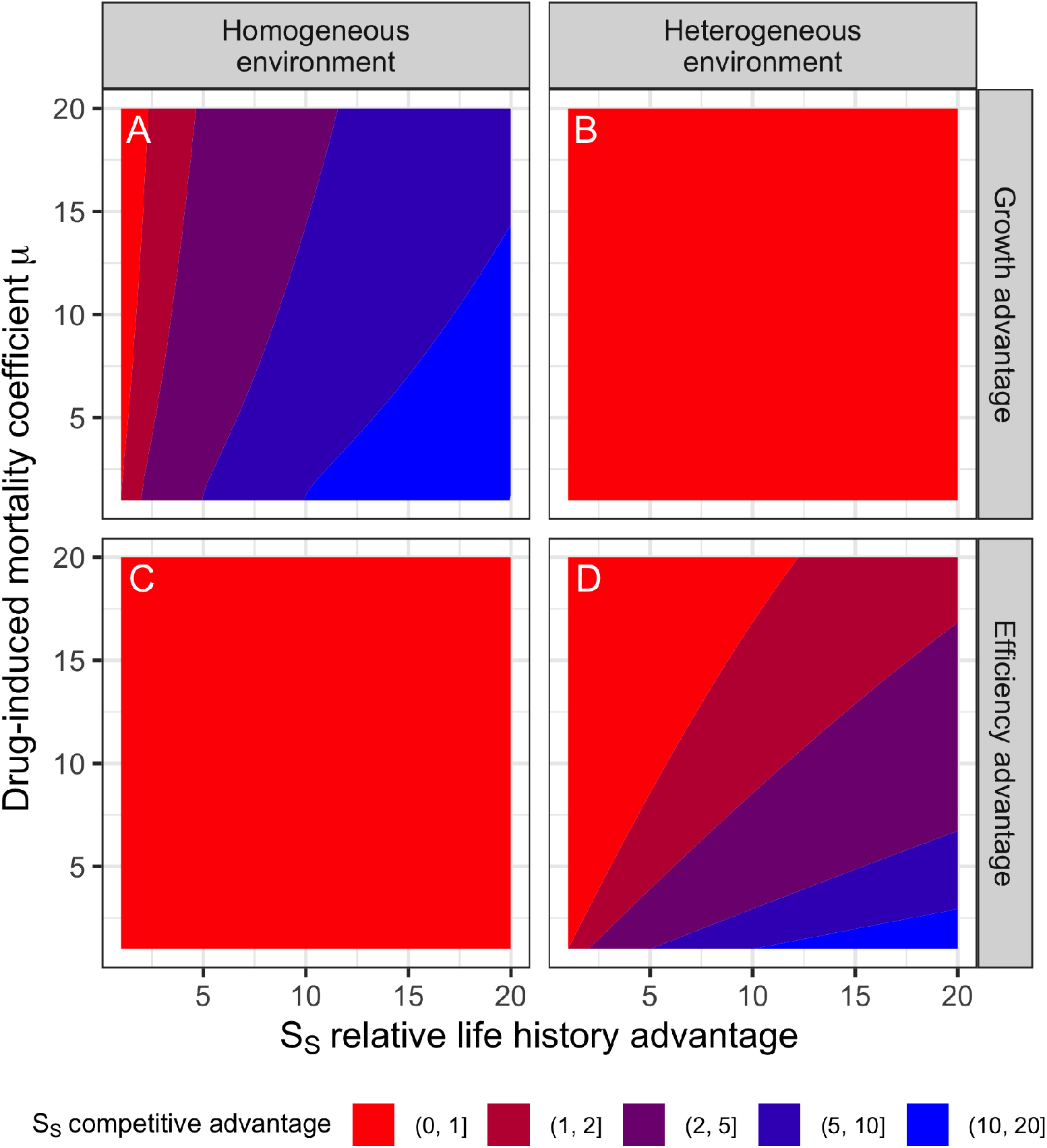
The expected competitive advantage for the drug-susceptible strain in homogeneous and heterogeneous environments as the life history advantage of the drug-sensitive strain varies (either growth rate or growth efficiency) and as the drug-induced mortality rate varies. The competitive advantage in a homogeneous environment is estimated by the ratio of the equilibrium resource levels maintained by each strain in a single patch 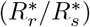: if 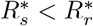, the susceptible strain suppresses resources more than the resistant strain, and we would predict it will outcompete the resistant strain in a homogeneous environment. The competitive advantage in a homogeneous environment is estimated by the ratio of the abundance equilibria each strain achieves in a single patch when it is alone 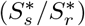:if 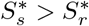, the susceptible strain has a higher abundance and we would predict it would have a colonization advantage over the resistant strain that will allow it to outcompete it in a heterogeneous environment. Panel (A) shows the competitive advantage of a fast-growing susceptible strain over a drug-resistant strain in a homogeneous environment. Panel (B) shows the (lack of) competitive advantage of a fast-growing susceptible strain over a drug-resistant strain in a heterogeneous environment. Panel (C) shows the (lack of) competitive advantage of an efficient susceptible strain over a drug-resistant strain in a homogeneous environment. Panel (D) shows the competitive advantage of an efficient susceptible strain over a drug-resistant strain in a heterogeneous environment.

Fig. 1A predicts a competitive advantage for the drug-sensitive strain as its growth advantage increases, whereas Fig. 1B predicts that the drug-resistant strain will always have a colonization advantage. Thus, we would predict that faster-growing drug-sensitive strains will outcompete resistant strains in homogeneous, but not heterogeneous, environments. Notably, these predictions reverse for drug-sensitive strains with an efficiency advantage. Fig. 1C predicts that the drug-resistant strain will always have a competitive advantage, whereas Fig. 1D predicts a colonization advantage for the drug-sensitive strain as its efficiency increases. Thus, we would predict that more efficient drug-sensitive strains will outcompete resistant strains in a heterogeneous, but not homogeneous, environment.

## Methods Part 2: Stochastic competition simulations

To assess these insights, we extend the consumer–resource model to a spatially structured environment and examine its dynamics using stochastic simulations. A stochastic model allows us to characterize the absolute abundance of each strain across all patches, rather than just the fraction of patches occupied by each strain, as in classic theory. In any deterministic model tracking strain abundances, once the strain with the lower single-patch 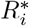reaches each patch, it will eventually exclude all other strains. Stochasticity allows for the random extinction and creation of patches and for random variation in how quickly each strain colonizes new patches.

We consider a landscape composed of multiple habitat patches connected by dispersal, generating variation in the persistence of local populations of simple bacterial communities and colonization opportunities. Each habitat patch follows the local consumer–resource dynamics described above, while individuals disperse between neighboring patches at a fixed colonization rate. These patches experience stochastic extinction events that remove all local populations and reset resource availability.

We computed the stochastic dynamics of each strain in a 100-patch spatially structured population with local migration (meaning that a migrating individual can only move to a neighboring patch; patches are assumed to be arrayed in a ring). For each scenario described above (varying relative growth rate, *γ*; relative growth efficiency, *ξ*; and drug-induced mortality coefficient, *µ*) we simulate the competition model 100 times, and then plot the prevalence of the drug-sensitive strain at the final time point, where prevalence is calculated as the fraction of the total bacterial population that is drug-sensitive across all patches. To generate a gradient of environmental heterogeneity, we vary the patch extinction rates, allowing us to evaluate how spatial structure alters the competitive balance predicted by single-patch theory.

## Results Part 2: Stochastic competition simulations

### Under what environmental conditions can competition suppress drug-resistance when resistant strains are already present?

Here, competitive outcomes reflect ecological interactions rather than evolutionary change *per se*, in that we assume that the drug-resistant strain is already present and at its evolutionarily stable drug-resistance strategy, 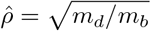Later, we relax this assumption and allow resistance to evolve dynamically from an initially drug-sensitive population. As above, we compare two scenarios, either the drug-sensitive strain is faster growing or more efficient relative to the drug-resistant strain (Fig. 2).

**Fig 2.**
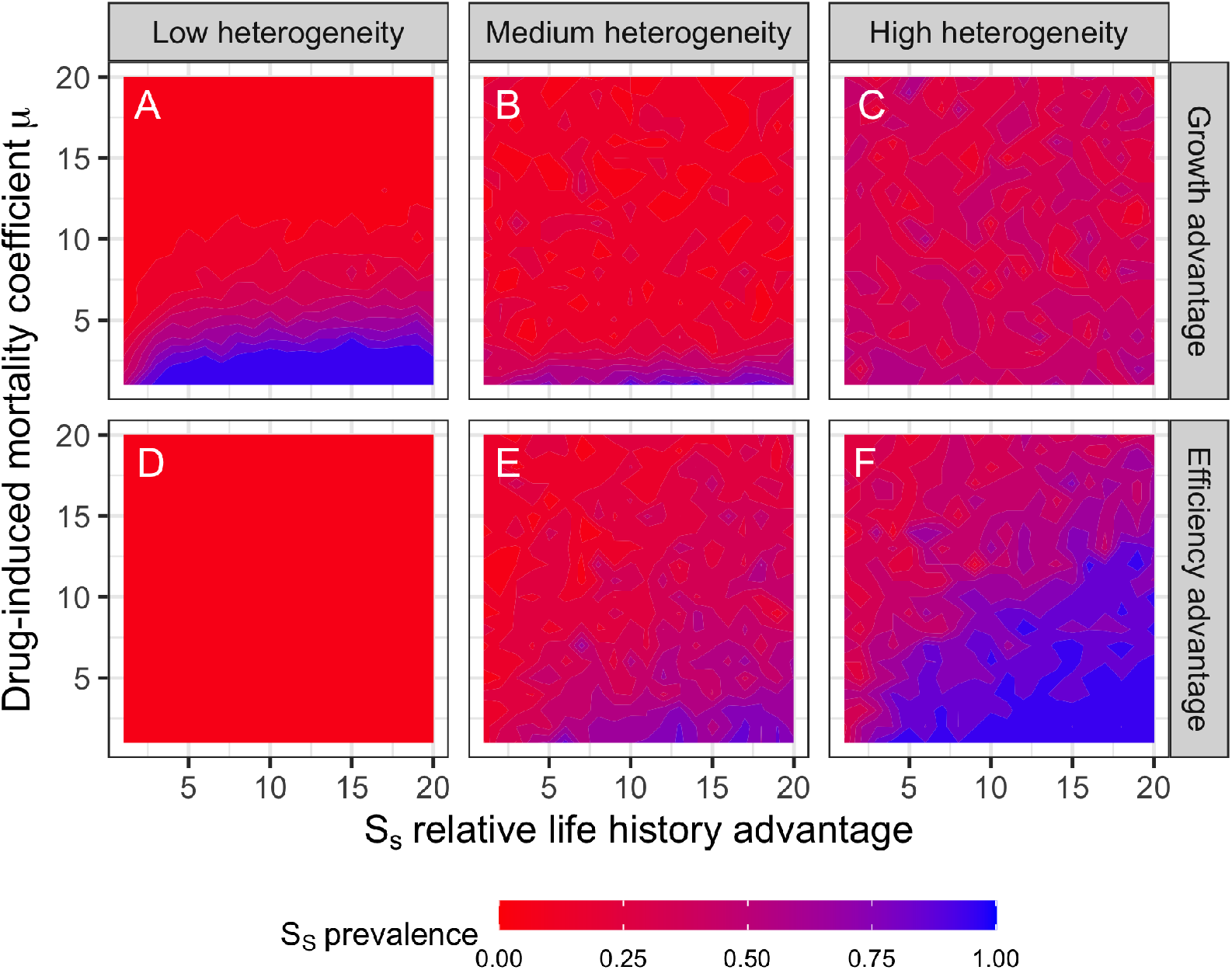
Prevalence of a drug-sensitive strain with either a growth (panels A-C) or efficiency (panels D-F) advantage over a drug-resistant competitor, when competition plays out in increasingly heterogeneous environments. Environmental heterogeneity is controlled by varying the patch extinction rate (low = 0.001, medium = 0.005, high = 0.01).

### Faster-growing drug-sensitive strains can only outcompete drug-resistant strains in homogeneous environments

In relatively homogeneous environments (low patch extinction rates), the drug-sensitive strain competitively excludes the faster-growing strain only when drug-induced mortality rate is low. Compared to Fig. 1A, we see that even the small amount of environmental heterogeneity introduced by spatial structure substantially reduces the parameter space in which the fast-growing drug-sensitive strain has an advantage (compare the size of the blue areas of Fig. 1A and Fig. 2A).

With moderate environmental heterogeneity (Fig. 2B), the competitive advantage of the resistant strain increases across most of the parameter space. At the highest level of heterogeneity (extinction rate), the outcome is largely governed by stochastic extinction, as evidenced by the mottled pattern in Fig. 2C that suggests modifying either the drug-induced mortality rate or growth advantage has no consistently predictable affect on the outcome of competition. However, the increasingly blue pattern (compared to Fig. 2A) reveals an advantage to the fast-growing drug-sensitive strain in this environment, because its fast growth makes it less sensitive to stochastic extinction.

However, this outcome results from stochasticity rather than from a change in the underlying competitive ordering. At the extreme levels of heterogeneity (extinction rate),rates, demographic stochasticity dominates and competitive outcomes become effectively random.

### More efficient, slower-growing drug-sensitive strains can outcompete drug-resistant strains in heterogeneous, low-resource environments

In relatively homogeneous environments (low patch extinction rates), the drug-resistant strain competitively excludes the more efficient but slower-growing drug-sensitive strain everywhere (Fig. 2D). This outcome is consistent with analytical predictions for homogeneous systems, where competition is governed by equilibrium resource requirements (*R*^*∗*^); under these conditions, efficiency does not affect competitive outcomes and confers no advantage (see Supplementary Information).

As environmental heterogeneity increases, however, competitive outcomes shift in a qualitatively different manner than in the first scenario.At intermediate and high levels of environmental heterogeneity, highly efficient drug-sensitive strains are predicted to exclude drug-resistant strains except under a narrow set of parameter values (Fig. 2E-F). Unlike the faster-growing competitor, the efficient strain gains a systematic advantage in heterogeneous environments because spatial structure converts differences in equilibrium population density (*S*^*∗*^) into colonization advantages.

Efficiency-based suppression depends on high equilibrium population densities ((*S*^*∗*^) to generate colonization advantages, which in turn depend on resource supply. Moreover, there is good empirical and theoretical evidence that efficiency is a K-selected strategy that is optimal in low resource environments, whereas fast growth is an r-selected strategy that is optimal in high resource environments (Raubenheimer and Simpson 1996, Marshall et al. 2023). To assess the robustness of efficiency-based suppression, we repeated this analysis under reduced resource availability.

Indeed, reducing resource availability enhances the efficiency-based competitive suppression observed under higher resource influx. In particular, in more homogeneous environments, the competitive advantage of the drug-resistant strain is weakened, as the efficient strain is more likely to persist and even outcompete the drug-resistant strain, especially when drug-induced mortality is low and efficiency is high. Similarly, the drug-sensitive strain is more likely to outcompete the drug-resistant strain under moderate levels of heterogeneity (patch extinction rates), even when the drug-induced mortality rate is high and efficiency is low. The results at the highest level of environmental heterogeneity (extinction rates) are fairly similar between the two resource levels, likely because stochasticity plays a larger role.

Together, these results suggest that when drug-resistant strains are already present in the environment, introducing a faster-growing drug-sensitive strain is unlikely to lead to reliable competitive suppression. Faster growth of the drug sensitive strain provides an advantage only under relatively homogeneous conditions. In contrast, spatial heterogeneity enables a distinct, efficiency-based suppression mechanism, whereby differences in equilibrium population density (*S*^*∗*^) translate into colonization advantages that allow efficient drug-sensitive strains to outcompete resistant strains. This competitive suppression is enhanced in low resource environments. These results indicate that increasing environmental heterogeneity could reduce the prevalence of drug-resistant strains.

We next extend this analysis to allow resistance to evolve dynamically and ask how environmental structure and competition jointly shape the emergence of drug-resistance.

### Methods Part 3: Stochastic evolution simulations

In the above section, we assumed that the drug-resistant strain had a resistance level set by the homogeneous environment ESS 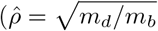 This potentially gives the largest possible advantage to the drug-resistant strain, which could bias our results if, for example, fast-growing strains are better at preventing the emergence of drug-resistance whereas efficient strains are better at competitively limiting emerged drug-resistant strains.

Here we assumed that all individuals of the drug-resistant strain were initially fully drug-sensitive (*ρ* = 1). When an individual of the drug-resistant strain replicated, the drug-resistance trait of its daughter cell was drawn from a lognormal distribution with mean equal to the individual’s trait and variance equal to *σ*^2^ = 0.01. Note that varying the value of *σ*^2^ would allow for faster or slower rates of evolution. This allows resistance evolution to emerge out of the ecological interactions between the drug-resistant and drug-sensitive strains, rather than specifying an explicit model of trait evolution [52].

## Results Part 3: Stochastic evolution simulations

### Spatial structure and resource limitation expand opportunities for efficiency-based competition to prevent the emergence of drug-resistance

Across a wide range of conditions, patterns of resistance emergence closely mirror the competitive outcomes observed when resistance is fixed (comparing Figs. 2 and 4).

Under relatively homogeneous environments, a faster-growing sensitive strain is better able to prevent the evolution of drug-resistance than an efficient sensitive strain (Fig. 4A vs. 4D). However, when the drug-induced mortality rate is very high, drug-resistance is very likely to emerge regardless of the life history advantage of the competitor.

Although a fast-growing strain is more likely to prevent the evolution of drug-resistance than to outcompete existing resistant strains in more heterogeneous environments (compare Fig. 2B, C to Fig. 4B, C), an efficient strain is still much more reliable at preventing the emergence of resistance (compare Fig. 4B, C to Fig. 4E, F). Indeed, especially at moderate levels of environmental heterogeneity (patch extinction rates), the efficient strain is better at preventing emergence than it would be at out competing an existing resistant strain (Fig. 2E vs. Fig. 4E). At the highest levels of environmental heterogeneity (patch extinction rates), however, the results for resistance competition and resistance emergence are very similar, again emphasizing the role of environmental stochasticity in mediating the outcome of competition (Fig. 2F vs. Fig. 4F).

Reducing resource availability modifies the outcome shown in Fig. 4 in a manner consistent with the fixed-resistance results (Fig. 2 vs. Fig. 3), but with an even stronger effect. In particular, under lower resource influx, efficiency-based prevention of resistance emergence is effective across a wider range of environmental heterogeneity. For example, when resources are low, a drug-sensitive efficient strain is better at preventing emergence of drug-resistance than a fast-growing strain in a high resource environment (compare Fig. 4A and 5A). However, at the highest levels of environmental heterogeneity (patch extinction rates), there is again an obvious influence of environmental stochasticity in allowing the resistant strain to potentially emerge.

**Fig 3.**
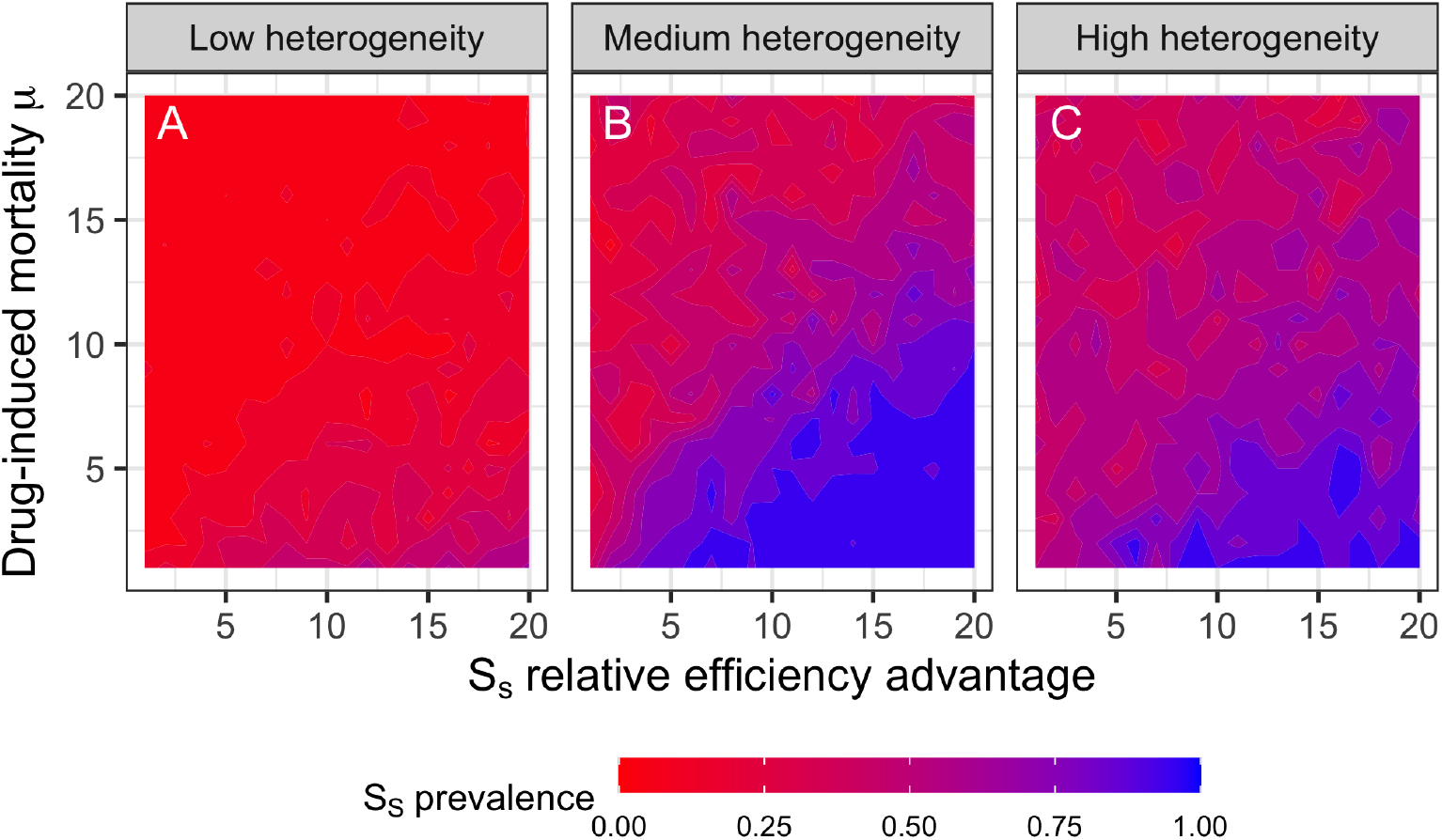
Prevalence of a drug-sensitive strain with an efficiency advantage over a drug-resistant competitor in resource-poor environments that vary in their heterogeneity (compare Fig. 3A-C against Fig. 2D-F).

**Fig 4.**
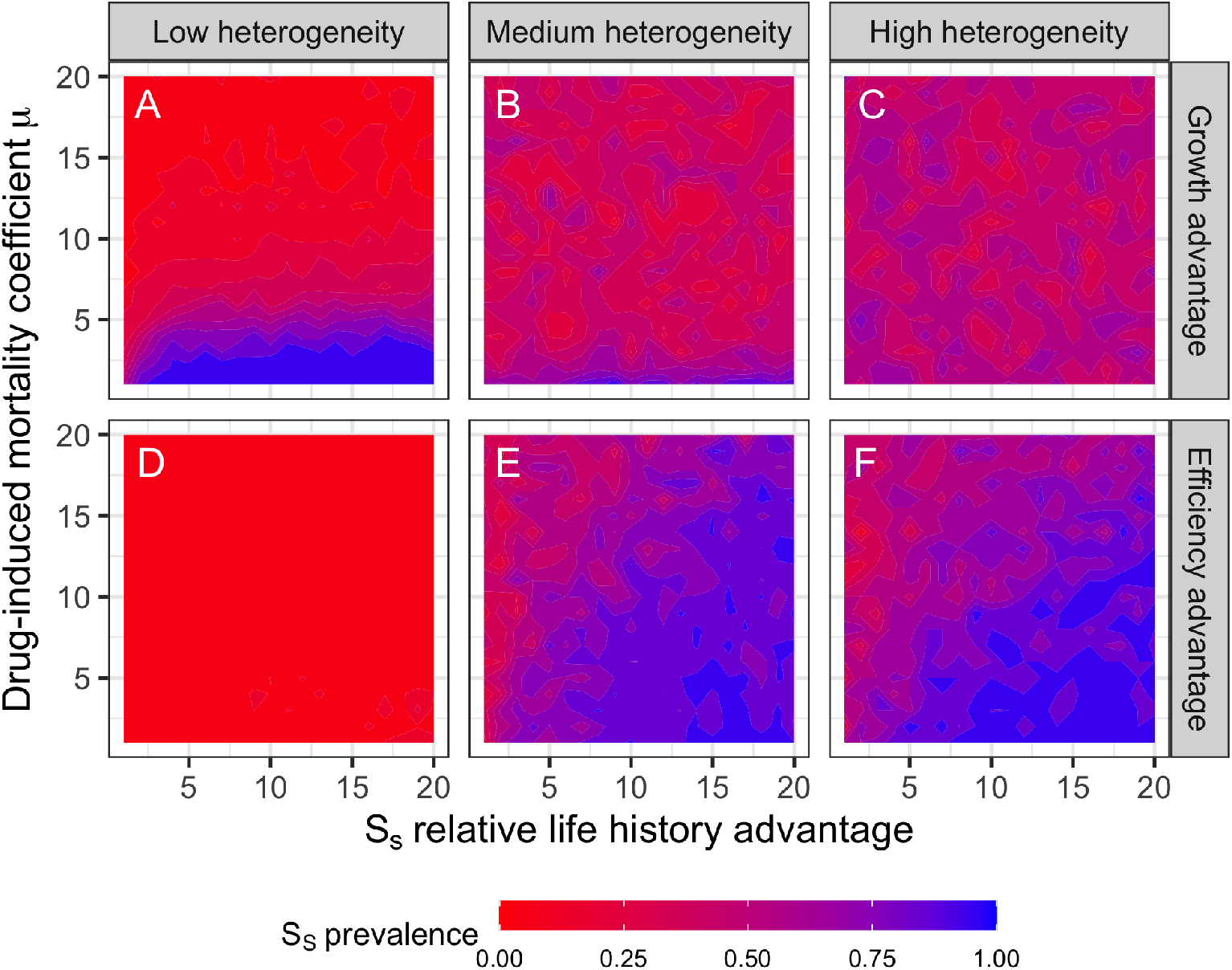
Prevalence of a drug-sensitive strain with either a growth (panels A-C) or efficiency (panels D-F) advantage over an evolving drug-resistant competitor, when competition plays out in increasingly heterogeneous environments. Environmental heterogeneity is controlled by varying the patch extinction rate (low = 0.001, medium = 0.005, high = 0.01). Rather than assuming that the drug-resistant strain has a resistance level equal to the ESS resistance, we assume that this strain is initially drug-sensitive but can evolve resistance under the growth–resistance trade-off.

**Fig 5.**
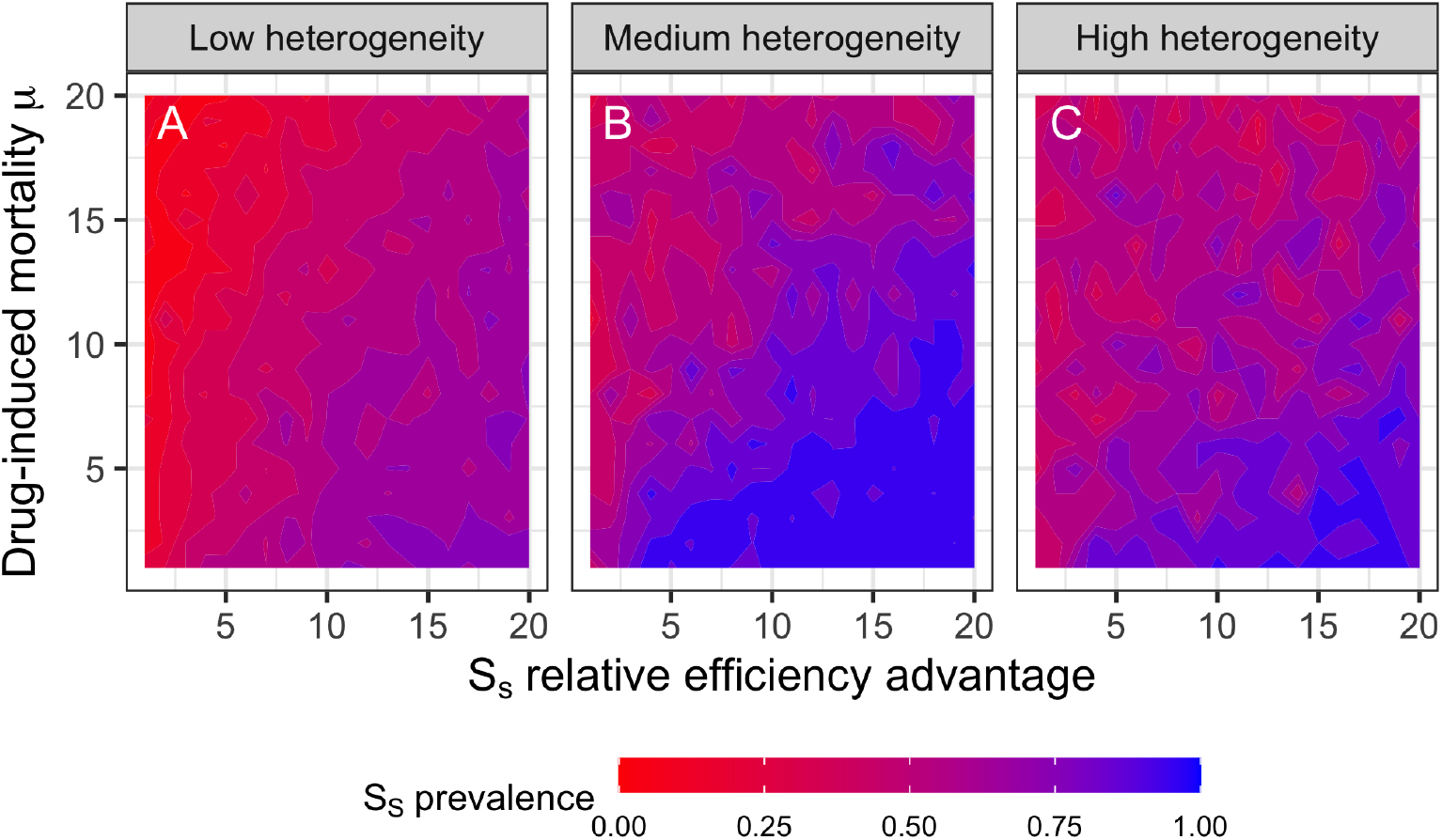
Prevalence of a drug-sensitive strain with an efficiency advantage over an evolving drug-resistant strain, under low resource availability, in increasingly heterogenous environments.

Together, these results demonstrate that allowing resistance to evolve does not alter the underlying ecological mechanisms that govern competition. Instead, resistance emerges or is prevented according to whether environmental heterogeneity enables efficiency-based, *S*^*∗*^-mediated suppression to operate.

## Discussion

Despite hundreds of thousands of drug-resistant microbes circulating across clinical, agricultural, and natural microbial communities, “hotspots” of antimicrobial resistance (AMR) are — all things considered — relatively rare. Why? Although we know many of the molecular and genetic factors that promote AMR, we know substantially less about the ecological factors that limiting the evolution of AMR [4, 8, 12, 55–57]. Our results suggest that a key factor is the way in which environmental conditions shape competitive interactions between drug-sensitive and drug-resistant strains.

Environmental heterogeneity does not simply strengthen or weaken competitive suppression; it changes the ecological mechanisms through which competition operates. This shift expands opportunities for drug-sensitive strains to suppress resistant competitors, with suppression strongest in heterogeneous environments with limited resources. These patterns are consistent regardless of whether AMR mutants are preexisting in the population (Figs. 2 and 3) or arise *de novo* (Figs. 4 and 5), suggesting that environmental structure can fundamentally alter the ecological conditions governing the evolution of AMR.

Consistent with previous theoretical and empirical work showing that drug-sensitive competitors can competitively suppress resistant lineages [58, 59], and that growth-rate advantages can counteract drug susceptibility [60], we find that faster-growing drug-sensitive strains can prevent resistance emergence under a restricted set of ecological conditions. However, this growth-based mechanism of competitive suppression operates under a relatively narrow range of conditions, specifically homogeneous environments with abundant resources and low drug-induced mortality. Under these conditions, competitive outcomes are governed by equilibrium resource requirements (*R*^*∗*^), and faster-growing drug-sensitive strains suppress resistance only when their growth advantage is sufficient to reduce its equilibrium resource requirement (*R*^*∗*^) below that of their resistant competitors. Beyond this region of parameter space, growth advantages alone are insufficient to suppress resistance.

In contrast, we also identify a second mechanism in which slower-growing but more resource-efficient competitors suppress resistance. This efficiency-based mechanism becomes increasingly important as environmental heterogeneity increases, particularly when resources are limiting. Under sufficient environmental heterogeneity, more efficient, slow growing drug-sensitive strains suppress drug-resistant competitors through a distinct mechanism in which differences in equilibrium population density (*S*^*∗*^) translate into colonization advantages: high efficiency leads to higher population densities, which leads to higher propagule pressure and thus, higher colonization success.

This efficiency-based competitive suppression operates across a broad region of parameter space when spatial turnover allows high-density patches to ‘seed’ recolonization. Lower resource availability further strengthens this mechanism. These findings are consistent with classic r-K tradeoffs in which efficient, slow growth (K-selected) strategies are favored under resource limitation, whereas inefficient, rapid growth (r-selection) is favored under resource abundance [61–63].

Although the model developed here is intentionally general, it relies on several simplifying assumptions. Most notably, we assumed a trade-off between resistance and growth rate. This assumption is supported by a substantial body of empirical work demonstrating growth-resistance trade-offs across diverse bacterial taxa [1, 45, 46]. For drug-sensitive competitors, we focused on growth rate and resource-use efficiency (yield) because they are fundamental components of microbial fitness widely linked with adaptation and evolution [45, 64]. While growth-rate costs of resistance have received considerable attention, few studies have examined how trade-offs involving resource-use efficiency shape resistance evolution [48–50]. Given the ubiquity of growth-efficiency trade-offs in microbial systems [45, 64], understanding how these traits interact with resistance may provide new insight into the ecological conditions that promote or suppress drug-resistance.

Additionally, drug-resistance can involve trade-offs in metabolism, resource acquisition, stress tolerance, and other physiological processes that are not fully captured by a simple growth-resistance trade-off [64]. For instance, metabolic adaptations often precede and facilitate the emergence of antimicrobial resistance and may alter additional traits other than growth rate and resource-use efficiency [48–50]. Consequently, resistance likely involves a more complex suite of trade-offs than previously appreciated, with important implications for evolutionary dynamics that are not explicitly captured in this model. The GEM modeling framework we developed here, however, can easily be extended to consider other trade-offs, as evolutionary dynamics emerge from the ecological model. Thus, exploring how alternative metabolic costs of resistance, or alternative pathways to resistance, are affected by competition only requires building those into the ecological model [51, 52].

Accounting for these additional trade-offs and competitive interactions could help expand opportunities for ecological management of AMR. For example, strategically manipulating competitive interactions offers a promising option to help manage the emergence and spread of drug-resistance in bacteria [15, 21], some cancers [16, 17, 31], and malaria [11, 18, 33, 65]. Our work helps address a critical question for designing such interventions: If the goal is to suppress resistance, would mitigation strategies that introduce fast-growing or resource-efficient drug-sensitive strains be more effective? Our results underscore that considering traits of the bacteria, spatial structure, and resource availability are crucial to addressing this question.

More broadly, an important implication of these results is that maintaining diversity in competitive strategies may increase opportunities for the suppression of resistant strains across a wide range of environmental conditions. The model developed here indicates that different strategies are favored under different ecological conditions: rapid growth is advantageous in resource-rich environments, whereas resource efficiency becomes increasingly important under resource limitation and spatial heterogeneity. Communities containing a diversity of competitive strategies may therefore retain multiple mechanisms for suppressing resistant strains as environmental conditions change [66].

Indeed, recent empirical evidence suggests that microbial diversity may play an underappreciated role in constraining or promoting antimicrobial resistance. For example, across soils, riverbeds, and human gut microbiomes, the abundance of antimicrobial resistance genes is negatively correlated with high microbial diversity [4, 9, 67]. However, the mechanisms underlying these associations remain poorly understood and an exciting area of active research [4, 57, 67].

On the one hand, higher microbial diversity could constrain resistance by increasing competitive interactions, diluting encounters between compatible hosts, and increasing the fitness costs of resistance [9]. Conversely, greater diversity could also promote resistance by increasing the number of potential bacterial hosts and interactions available for horizontal gene transfer [10]. Thus, whether microbial diversity constrains or promotes resistance likely depends on how community composition and ecological interactions jointly alter the balance between competition, transmission, and selection [66]. Our results suggest that environmental heterogeneity may be one important component of this balance because it determines which competitive strategies are favored and, consequently, which mechanisms of competitive suppression are most effective.

This perspective may help reconcile why previous theoretical and empirical studies examining the effects of spatial structure on resistance evolution have reached contrasting conclusions. Some studies conclude that spatial structure slows resistance evolution, whereas others find that it accelerates adaptation. Our results suggest that these different outcomes may arise, at least in part, because studies have focused on different sources of spatial heterogeneity. Spatial variation in antimicrobial exposure and resource availability need not favor the same competitive strategies and may therefore generate different opportunities for resistant strains to emerge and spread.

Much of the existing literature has focused on spatial variation in antimicrobial exposure itself and assumed that competing strains differ primarily in their resistance phenotypes, often involving a growth-resistance trade-off. In effect, these models assume heterogeneous antimicrobial concentrations and relatively homogeneous resource environments [39]. These models have primarily been developed to investigate the effects of spatial variation within treated hosts, where pharmacokinetic differences among tissues and organs generate spatial variation in antimicrobial concentrations.

Bacteria may also move among hosts, experiencing different antimicrobial, immune, and resource environments, further contributing to spatial variation in selection [19, 40–44].

Our model reverses these assumptions by allowing resource distributions to vary spatially while antimicrobial exposure remains homogeneous. Under these conditions, homogeneous environments favor fast-growing drug-sensitive strains, whereas heterogeneous environments favor more resource-efficient competitors. Whether heterogeneity arises from variation in antimicrobial exposure or resource availability, successful lineages must frequently colonize and spread into newly favorable environments. Despite these differences in assumptions, a common conclusion emerges from both our work and previous studies: spatial heterogeneity often couples adaptation with range expansion.

Antimicrobial resistance (AMR) represents a significant and increasing global threat to public health and agriculture and carries unknown consequences for natural ecosystems and wildlife. Increasing evidence suggests that anthropogenic environmental changes, beyond antimicrobial usage, are responsible for promoting the emergence and spread of drug-resistant microbes [4, 57, 67, 68]. However, identifying mechanistic links between environmental drivers and emergence remains exceptionally challenging and largely overlooked in the global response to combating AMR [5, 7, 8, 56]. This oversight arises, in part, because surprisingly little is known about how ecological interactions influence the evolution of antimicrobial resistance [4, 5, 8, 56]. Our results provide potential mechanisms linking anthropogenic changes to AMR via the effects of spatial structure and microbial traits on competitive interactions.

The main takeaway from the model developed here is not just that spatial structure or resource availability matter for AMR; the importance of these mechanisms has a long history in ecological and evolutionary theory [14, 22, 29, 30]. Instead, the central lesson is that homogenization of resources may help promote AMR. Because many human activities can homogenize environments (e.g., invasive plant species, chemically intensive monoculture agriculture typical across much of the US Corn Belt), it is critical to determine whether they do so at the spatial scales relevant to microbial competition [4, 68, 69]. Future studies integrating AMR surveillance data and environmental meta-data with eco-evolutionary models, such as the one developed here, could help move environmental heterogeneity and microbial community structure from correlates of resistance into mechanistic predictors of AMR.

## Supporting information

Analytical results and code for the main text

## Data Availability

All associated code necessary to reproduce the analysis reported here is available in the Supplementary Information.

## References

1. Haugan MS, Løbner-Olesen A, Frimodt-Møller N. Comparative activity of ceftriaxone, ciprofloxacin, and gentamicin as a function of bacterial growth rate probed by Escherichia coli chromosome replication in the mouse peritonitis model. Antimicrobial Agents and Chemotherapy. 2019;63(2):10–1128.

2. Deb LC, Jara M, Lanzas C. Early evaluation of the Food and Drug Administration (FDA) guidance on antimicrobial use in food animals on antimicrobial resistance trends reported by the National Antimicrobial Resistance Monitoring System (2012–2019). One Health. 2023;17:100580.

3. Steinberger AJ, Nickodem CA, Leite de Campos J, Kates AE, Goldberg TL, Safdar N, et al. Antimicrobial use contributes to resistance gene enrichment across cattle groups on commercial dairy farms. bioRxiv. 2026:2026–05.

4. Nickodem CA, Tran PQ, Neeno-Eckwall E, Congdon AG, Sanford GR, Silva EM, et al. Soil management strategies drive divergent impacts on pathogens and environmental resistomes. Scientific Reports. 2025;15(1):43215.

5. Woolhouse M, Ward M, Van Bunnik B, Farrar J. Antimicrobial resistance in humans, livestock and the wider environment. Philosophical Transactions of the Royal Society B: Biological Sciences. 2015;370(1670).

6. Dolgin E. Combating antibiotic resistance from the ground up. Proceedings of the National Academy of Sciences. 2016;113(42):11642–3.

7. Organization WH. Global antimicrobial resistance and use surveillance system (GLASS) report 2022. World Health Organization; 2022.

8. Bottery MJ, Pitchford JW, Friman VP. Ecology and evolution of antimicrobial resistance in bacterial communities. The ISME Journal. 2021;15(4):939–48.

9. Chen QL, An XL, Zheng BX, Gillings M, Peñuelas J, Cui L, et al. Loss of soil microbial diversity exacerbates spread of antibiotic resistance. Soil Ecology Letters. 2019;1(1):3–13.

10. Klümper U, Gionchetta G, Catão E, Bellanger X, Dielacher I, Elena AX, et al. Environmental microbiome diversity and stability is a barrier to antimicrobial resistance gene accumulation. Communications Biology. 2024;7(1):706.

11. Babiker HA, Hastings IM, Swedberg G. Impaired fitness of drug-resistant malaria parasites: evidence and implication on drug-deployment policies. Expert review of anti-infective therapy. 2009;7(5):581–93.

12. Andersson DI, Hughes D. Antibiotic resistance and its cost: is it possible to reverse resistance? Nature Reviews Microbiology. 2010;8(4):260–71.

13. Wargo AR, Huijben S, De Roode JC, Shepherd J, Read AF. Competitive release and facilitation of drug-resistant parasites after therapeutic chemotherapy in a rodent malaria model. Proceedings of the National Academy of Sciences. 2007;104(50):19914–9.

14. Read AF, Day T, Huijben S. The evolution of drug resistance and the curious orthodoxy of aggressive chemotherapy. Proceedings of the National Academy of Sciences. 2011;108(supplement 2):10871–7.

15. Day T, Huijben S, Read AF. Is selection relevant in the evolutionary emergence of drug resistance? Trends in microbiology. 2015;23(3):126–33.

16. Gatenby RA, Brown J, Vincent T. Lessons from applied ecology: cancer control using an evolutionary double bind. Cancer research. 2009;69(19):7499–502.

17. Colijn C, Cohen T. How competition governs whether moderate or aggressive treatment minimizes antibiotic resistance. Elife. 2015;4:e10559.

18. Huijben S, Bell AS, Sim DG, Tomasello D, Mideo N, Day T, et al. Aggressive chemotherapy and the selection of drug resistant pathogens. PLOS Pathogens. 2013;9(9):e1003578.

19. Baym M, Lieberman TD, Kelsic ED, Chait R, Gross R, Yelin I, et al. Spatiotemporal microbial evolution on antibiotic landscapes. Science. 2016;353(6304):1147–51.

20. Bacevic K, Noble R, Soffar A, Wael Ammar O, Boszonyik B, Prieto S, et al. Spatial competition constrains resistance to targeted cancer therapy. Nature Communications. 2017;8(1):1995.

21. Hansen E, Karslake J, Woods RJ, Read AF, Wood KB. Antibiotics can be used to contain drug-resistant bacteria by maintaining sufficiently large sensitive populations. PLOS Biology. 2020;18(5):e3000713.

22. Day T, Read AF. Does high-dose antimicrobial chemotherapy prevent the evolution of resistance? PLOS Computational Biology. 2016;12(1):e1004689.

23. Hsu PP, Sabatini DM. Cancer cell metabolism: Warburg and beyond. Cell. 2008;134(5):703–7.

24. Gillies RJ, Verduzco D, Gatenby RA. Evolutionary dynamics of carcinogenesis and why targeted therapy does not work. Nature Reviews Cancer. 2012;12(7):487–93.

25. Ross-Gillespie A, Weigert M, Brown SP, Kümmerli R. Gallium-mediated siderophore quenching as an evolutionarily robust antibacterial treatment. Evolution, medicine, and public health. 2014;2014(1):18–29.

26. Garber K. Targeting copper to treat breast cancer. American Association for the Advancement of Science; 2015.

27. Tilman D. Resource competition and community structure. 17. Princeton university press; 1982.

28. Tilman D. Competition and biodiversity in spatially structured habitats. Ecology. 1994;75(1):2–16.

29. Tilman EA, Tilman D, Crawley MJ, Johnston A. Biological weed control via nutrient competition: potassium limitation of dandelions. Ecological Applications. 1999;9(1):103–11.

30. Smith VH, Holt RD. Resource competition and within-host disease dynamics. Trends in ecology & evolution. 1996;11(9):386–9.

31. Enriquez-Navas PM, Kam Y, Das T, Hassan S, Silva A, Foroutan P, et al. Exploiting evolutionary principles to prolong tumor control in preclinical models of breast cancer. Science translational medicine. 2016;8(327):327ra24–4.

32. Wale N, Sim DG, Read AF. A nutrient mediates intraspecific competition between rodent malaria parasites in vivo. Proceedings of the Royal Society B: Biological Sciences. 2017;284(1859):20171067.

33. Wale N, Sim DG, Jones MJ, Salathe R, Day T, Read AF. Resource limitation prevents the emergence of drug resistance by intensifying within-host competition. Proceedings of the National Academy of Sciences. 2017;114(52):13774–9.

34. Wright S. Statistical theory of evolution. Journal of the American Statistical Association. 1931;26(173A):201–8.

35. Holt RD, Gomulkiewicz R. The evolution of species’ niches: a population dynamic perspective. Case studies in mathematical modelling: ecology, physiology, and cell biology. 1997:25–50.

36. May R, McLean AR. Theoretical ecology: principles and applications. OUP Oxford; 2007.

37. González-García I, Solé RV, Costa J. Metapopulation dynamics and spatial heterogeneity in cancer. Proceedings of the National Academy of Sciences. 2002;99(20):13085–9.

38. Cremer J, Melbinger A, Frey E. Growth dynamics and the evolution of cooperation in microbial populations. Scientific Reports. 2012;2(1):281.

39. Hernández-Navarro L, Distefano K, Täuber UC, Mobilia M. Slow spatial migration can help eradicate cooperative antimicrobial resistance in time-varying environments. PLOS Computational Biology. 2026;22(3):e1013997.

40. De Jong MG, Wood KB. Tuning spatial profiles of selection pressure to modulate the evolution of drug resistance. Physical review letters. 2018;120(23):238102.

41. Hermsen R, Hwa T. Sources and sinks: a stochastic model of evolution in heterogeneous environments. Physical review letters. 2010;105(24):248104.

42. Zhang Q, Lambert G, Liao D, Kim H, Robin K, Tung Ck, et al. Acceleration of emergence of bacterial antibiotic resistance in connected microenvironments. Science. 2011;333(6050):1764–7.

43. Akter M, Bailey SF. Agar concentration affects the evolution of antibiotic resistance and population genomics in experimental populations Pseudomonas aeruginosa. bioRxiv. 2024:2024–09.

44. France MT, Cornea A, Kehlet-Delgado H, Forney LJ. Spatial structure facilitates the accumulation and persistence of antibiotic-resistant mutants in biofilms. Evolutionary Applications. 2019;12(3):498–507.

45. Lipson DA. The complex relationship between microbial growth rate and yield and its implications for ecosystem processes. Frontiers in Microbiology. 2015;6:615.

46. Greulich P, Scott M, Evans MR, Allen RJ. Growth-dependent bacterial susceptibility to ribosome-targeting antibiotics. Molecular systems biology. 2015;11(3):MSB145949.

47. Frank SA. Microbial life history: The fundamental forces of biological design. Princeton University Press; 2022.

48. Lopatkin AJ, Bening SC, Manson AL, Stokes JM, Kohanski MA, Badran AH, et al. Clinically relevant mutations in core metabolic genes confer antibiotic resistance. Science. 2021;371(6531):eaba0862.

49. Lopatkin AJ, Stokes JM, Zheng EJ, Yang JH, Takahashi MK, You L, et al. Bacterial metabolic state more accurately predicts antibiotic lethality than growth rate. Nature Microbiology. 2019;4(12):2109–17.

50. Ahmad M, Aduru SV, Smith RP, Zhao Z, Lopatkin AJ. The role of bacterial metabolism in antimicrobial resistance. Nature Reviews Microbiology. 2025;23(7):439–54.

51. DeLong JP, Gibert JP. Gillespie eco-evolutionary models (GEMs) reveal the role of heritable trait variation in eco-evolutionary dynamics. Ecology and evolution. 2016;6(4):935–45.

52. DeLong JP, Cressler CE. Stochasticity directs adaptive evolution toward nonequilibrium evolutionary attractors. Ecology. 2023;104(1):e3873.

53. Doebeli M, Ispolatov Y, Simon B. Towards a mechanistic foundation of evolutionary theory. Elife. 2017;6:e23804.

54. Pfeiffer T, Schuster S, Bonhoeffer S. Cooperation and competition in the evolution of ATP-producing pathways. Science. 2001;292(5516):504–7.

55. Pradier L, Bedhomme S. Ecology, more than antibiotics consumption, is the major predictor for the global distribution of aminoglycoside-modifying enzymes. Elife. 2023;12:e77015.

56. Berini III J, Fouilloux CA, Neeno-Eckwall E, Alexander H, Choi E, Vaziri G, et al. A multi-scale ecological approach to assessing antimicrobial resistance in a freshwater fish. bioRxiv. 2026:2026–05.

57. Kalanxhi E, Laxminarayan R. Climate change and antimicrobial resistance. Nature Reviews Microbiology. 2026:1–12.

58. Hansen E, Woods RJ, Read AF. How to use a chemotherapeutic agent when resistance to it threatens the patient. PLOS biology. 2017;15(2):e2001110.

59. Czuppon P, Day T, Débarre F, Blanquart F. A stochastic analysis of the interplay between antibiotic dose, mode of action, and bacterial competition in the evolution of antibiotic resistance. PLOS Computational Biology. 2023;19(8):e1011364.

60. Amor DR, Gore J. Fast growth can counteract antibiotic susceptibility in shaping microbial community resilience to antibiotics. Proceedings of the National Academy of Sciences. 2022;119(15):e2116954119.

61. Raubenheimer D, Simpson SJ. Integrating nutrition: a geometrical approach. Entomologia experimentalis et applicata. 1999;91(1):67–82.

62. Simpson S, Raubenheimer D. The geometric analysis of feeding and nutrition: a user’s guide. Journal of Insect Physiology. 1995;41(7):545–53.

63. Marshall DJ, Cameron HE, Loreau M. Relationships between intrinsic population growth rate, carrying capacity and metabolism in microbial populations. The ISME Journal.

64. Cheng C, O’brien EJ, McCloskey D, Utrilla J, Olson C, LaCroix RA, et al. Laboratory evolution reveals a two-dimensional rate-yield tradeoff in microbial metabolism. PLOS Computational Biology. 2019;15(6):e1007066.

65. Pollitt LC, Huijben S, Sim DG, Salathé RM, Jones MJ, Read AF. Rapid response to selection, competitive release and increased transmission potential of artesunate-selected Plasmodium chabaudi malaria parasites. PLOS Pathogens. 2014;10(4):e1004019.

66. Patangia DV, Anthony Ryan C, Dempsey E, Paul Ross R, Stanton C. Impact of antibiotics on the human microbiome and consequences for host health. Microbiologyopen. 2022;11(1):e1260.

67. Bagra K, Bellanger X, Merlin C, Singh G, Berendonk TU, Klümper U. Environmental stress increases the invasion success of antimicrobial resistant bacteria in river microbial communities. Science of the Total Environment. 2023;904:166661.

68. Kelbrick M, Hesse E, O’Brien S. Cultivating antimicrobial resistance: how intensive agriculture ploughs the way for antibiotic resistance. Microbiology. 2023;169(8):001384.

69. Groffman PM, Cavender-Bares J, Bettez ND, Grove JM, Hall SJ, Heffernan JB, et al. Ecological homogenization of urban USA. Frontiers in Ecology and the Environment. 2014;12(1):74–81.

