## Supplementary material for "Efficiency is better than speed for competitive suppression of drug-resistance": Analytical results and code for the main text

### Adaptive Dynamics Analysis of a Homogeneous Environment

In a spatially homogeneous environment, the dynamics of two strains of bacteria, $S_s$ (a drug-susceptible strain) and $S_r$ (a drug-resistant strain), competing for a single limiting resource are determined by the simple model:

$$\begin{aligned}
\frac{dR}{dt} &= i - oR - \frac{1}{Y_s}\frac{g_s R}{h_s + R}S_s - \frac{1}{Y_r}\frac{g_r(\rho)R}{h_r + R}S_r \\
\frac{dS_s}{dt} &= \frac{g_s R}{h_s + R}S_s - (m_b + m_d)\,S_s \\
\frac{dS_r}{dt} &= \frac{g_r(\rho)R}{h_r + R}S_r - (m_b + m_d(\rho))\,S_r
\end{aligned}$$

The two strains are assumed to uptake resources according to Michaelis-Menten kinetics. The resource ($R$) is assumed to follow semi-chemostat dynamics, with a constant resource influx at rate $i$ and removal at rate $o$. Both strains uptake resource assuming Michaelis-Menten (Monod) uptake kinetics, with maximum growth rates $g_{s,r}$ and half-saturation constants $h_{s,r}$. The yield parameters $Y_{s,r}$ quantify the bacterial biomass produced per unit of resource biomass; although $Y_{s,r}$ is typically defined as the yield, it can be more intuitive to think of it as quantifying growth efficiency, with higher values of $Y$ implying more efficient biomass production (Frank 2022). Bacterial biomass is lost at a background rate $m_b$ and a drug-induced mortality rate $m_d$.

The resistant strain, $S_r$, has a trait, $\rho$ that determines both its growth rate ($g_r(\rho)$) and its drug-induced mortality rate ($m_d(\rho)$). We assume the following following functional relationships:

$$g_r(\rho) = \frac{g}{\rho}, \qquad m_d(\rho) = \frac{m_d}{\rho^2}.$$

These functional forms capture a growth–resistance trade-off in which increasing resistance (higher $\rho$) reduces growth while lowering susceptibility to antimicrobials (drug-induced mortality). These functional forms were chosen because, in a spatially homogeneous environment, this trade-off gives rise to a single evolutionarily stable resistance strategy, $\hat{\rho}$:

$$\hat{\rho} = \sqrt{\frac{m_d}{m_b}}.$$

To see how we arrive at this conclusion, we use an adaptive dynamics approach. Specifically, we consider whether a mutant strain, $S_m$, with resistance trait, $\rho_m$, can invade a resident population of resistant bacteria, $S_r$ with resistance trait, $\rho_s$:

$$\begin{aligned}
\frac{dR}{dt} &= i - oR - \frac{1}{Y_r}\frac{g_r(\rho_r)R}{h_s + R}S_r - \frac{1}{Y_r}\frac{g_r(\rho_m)R}{h_r + R}S_m, \\
\frac{dS_r}{dt} &= \frac{g_r(\rho_r)R}{h_r + R}S_r - (m_b + m_d(\rho_r))\,S_r, \\
\frac{dS_m}{dt} &= \frac{g_r(\rho_m)R}{h_r + R}S_m - (m_b + m_d(\rho_m))\,S_m.
\end{aligned}$$

The mutant strain can invade if its per-capita growth rate at the equilibrium resource level set by the resident strain is positive, that is, if

$$F = \frac{g_r(\rho_m)\hat{R}}{h_r + \hat{R}} - (m_b + m_d(\rho_m)) > 0,$$

where

$$\hat{R} = \frac{h_r\,(m_b + m_d(\rho_r))}{g_r(\rho_r) - (m_b + m_d(\rho_r))}$$

is the equilibrium resource level when $S_r$ is interacting with resources alone.

Note that we can rearrange $F > 0$ to gain more ecological insight. Specifically, if we solve the inequality for $\hat{R}$, we find the following expression:

$$\frac{h_r\,(m_b + m_d(\rho_m))}{g_r(\rho_m) - (m_b + m_d(\rho_m))} < \hat{R}.$$

Notice that the expression on the left is just the equilibrium resource level when $S_m$ is interacting with resources alone. Thus, the mutant can invade if it suppresses resources to a lower level than the resident. This confirms the intuition that competition minimizes the equilibrium resource abundance.

To find potential evolutionarily stable resistance strategies, we first plug our expression for $\hat{R}$ into the invasion condition $F$; we then differentiate the invasion condition with respect to the evolving trait $\rho_m$; we then set $\rho_m = \rho_r = \hat{\rho}$ and simplify; finally, we set this condition equal to zero and solve for $\hat{\rho}$. Following this process leads to the following – any evolutionarily stable strategy $\rho_r$ must satisfy this condition:

$$\left[\frac{\partial F}{\partial \rho_m}\right]_{\rho_m=\rho_r=\hat{\rho}} = \frac{(m_b + m_d(\rho))\,g_r'(\rho)}{g_r(\rho)} - m_d'(\rho) = 0.$$

Plugging $g_r(\rho) = g/\rho$ and $m_d(\rho) = m_d/\rho^2$ into leads to

$$\frac{m_d - m_b\hat{\rho}^2}{\hat{\rho}^3} = 0,$$

which simplifies to $\hat{\rho} = \sqrt{\frac{m_d}{m_b}}$.

To confirm that $\hat{\rho}$ is, in fact, an evolutionarily stable strategy, we need to confirm that it is a maximum by taking the second derivative of our invasion condition $F$ with respect to $\rho_m$ and set $\rho_m = \rho_r = \hat{\rho}$ and confirming that the resulting expression is negative. This yields:

$$\left[\frac{\partial^2 F}{\partial \rho_m^2}\right]_{\rho_m=\rho_r=\hat{\rho}} = -\frac{2m_b^2}{m_d}.$$

Since this expression is always negative, we know that any solution $\hat{\rho}$ will be evolutionarily stable.

Finally, to confirm that $\hat{\rho}$ is also convergence stable, we take the derivative of $F$ with respect to $\rho_m$, set $\rho_m = \rho_r$ and take the derivative with respect to $\rho_r$, and then set $\rho_r = \hat{r}$ and confirm that the resulting expression is negative. This yields:

$$\left[\frac{\partial}{\partial \rho_r}\left[\frac{\partial R}{\partial \rho_m}\right]_{\rho_m=\rho_r}\right]_{\rho_r=\hat{\rho}} = -\frac{2m_b^2}{m_d}.$$

This verifies that $\hat{\rho} = \sqrt{\frac{m_d}{m_b}}$ is, in fact, a fitness-maximizing strategy that can be reached by gradual evolutionary steps.

### Code to Reproduce Results

Figure 1:

```r
## Figure 1
calc_rel_equil_1 <- function(mort_coef, growth_coef) {
  Yr=0.1
  Ys=0.1
  gr=5
  gs=5*growth_coef
  h=20
  m0=0.01
  m1=0.01*mort_coef
  i=100
  o=1

  ## ESS resistance
  r = sqrt(m1/m0)

  ## Resource equilibria for each strain
  Rr = h*(m0+m1/r^2)/(gr/r-m0-m1/r^2)
  Rs = h*(m0+m1)/(gs-m0-m1)

  ## Density equilibria for each strain
  Sr = (i-o*Rr)*Yr*(h+Rr)/(gr/r*Rr)
  Ss = (i-o*Rs)*Ys*(h+Rs)/(gs*Rs)

  return(c(Rr/Rs, Ss/Sr))
}

calc_rel_equil_2 <- function(mort_coef, yield_coef) {
  Yr=0.1
  Ys=0.1*yield_coef
  gr=5
  gs=0.5
  h=20
  m0=0.01
  m1=0.01*mort_coef
  i=100
  o=1

  ## ESS resistance
  r = sqrt(m1/m0)

  ## Resource equilibria for each strain
  Rr = h*(m0+m1/r^2)/(gr/r-m0-m1/r^2)
  Rs = h*(m0+m1)/(gs-m0-m1)

  ## Density equilibria for each strain
  Sr = (i-o*Rr)*Yr*(h+Rr)/(gr/r*Rr)
  Ss = (i-o*Rs)*Ys*(h+Rs)/(gs*Rs)

  return(c(Rr/Rs, Ss/Sr))
}

equil <- expand.grid(mort_coef=seq(1,20,0.1), growth_coef=seq(1,20,0.1))
sapply(1:nrow(equil), function(i) calc_rel_equil_1(equil[i,1], equil[i,2])) %>%
```

```
  t %>%
  as.data.frame() %>%
  mutate(mort_coef=equil$mort_coef,
         Ss_coef=equil$growth_coef,
         Ss_advantage="Growth advantage") -> rel_equil_1
colnames(rel_equil_1)[1:2] <- c("Homogeneous\nenvironment","Heterogeneous\nenvironment")

equil <- expand.grid(mort_coef=seq(1,20,0.1), yield_coef=seq(1,20,0.1))
sapply(1:nrow(equil), function(i) calc_rel_equil_2(equil[i,1], equil[i,2])) %>%
  t %>%
  as.data.frame() %>%
  mutate(mort_coef=equil$mort_coef,
         Ss_coef=equil$yield_coef,
         Ss_advantage="Efficiency advantage") -> rel_equil_2
colnames(rel_equil_2)[1:2] <- c("Homogeneous\nenvironment","Heterogeneous\nenvironment")

rbind(rel_equil_1, rel_equil_2) %>%
  pivot_longer(., cols=1:2) -> rel_equil
rel_equil$Ss_advantage = factor(rel_equil$Ss_advantage, levels=c("Growth advantage", "Efficiency advanta
rel_equil$name = factor(rel_equil$name, levels=c("Homogeneous\nenvironment", "Heterogeneous\nenvironmen

ann_df = expand.grid(name=levels(rel_equil$name),
                      Ss_advantage = levels(rel_equil$Ss_advantage)
                      )
ann_df$label = LETTERS[1:4]
ann_df$x = -Inf
ann_df$y = Inf

png(file="Fig1_homogeneous_environment_predictions.png", height=5, width=4.5, units='in', res=400)
rel_equil %>%
  ggplot(., aes(x=Ss_coef, y=mort_coef, z=value)) +
  geom_contour_filled(breaks=c(0,1,2,5,10,20)) +
  scale_fill_manual(values=colorRampPalette(c("red","blue"))(5)) +
  geom_text(
    data = ann_df,
    aes(x = x, y = y, label = label),
    hjust = -1,
    vjust = 2,
    color="white",
    inherit.aes = FALSE,
    size = 4
  ) +
  facet_grid(Ss_advantage~name) +
  xlab(expression(S[S] ~ "relative life history advantage")) +
  ylab(expression("Drug-induced mortality coefficient" ~ mu)) +
  labs(fill=expression(S[S] ~ "competitive advantage")) +
  theme_bw() +
  theme(legend.position="bottom",
        legend.key.size=unit(0.5,"cm"),
        legend.text=element_text(size=7),
        legend.title=element_text(size=8),
        legend.spacing.y=unit(2,"mm"),
        legend.margin=margin(1,1,1,1))
```

```r
dev.off()
```

C++ code to simulate competition between two strains that differ in their life histories:

```cpp
#include <Rcpp.h>
using namespace Rcpp;

// [[Rcpp::export]]
List stoch_spatial_comp_3(NumericVector params) {
  // Extract all relevant model parameters and algorithm parameters from params
  // Note that it is critical that parameters are specified in EXACTLY the order they are extracted in
  // Extract bacteria parameter values from params
  double Y1 = params[0]; // strain 1 yield
  double Y2 = params[1]; // strain 2 yield
  double g1 = params[2]; // strain 1 maximum growth rate
  double g2 = params[3]; // strain 2 maximum growth rate
  double h1 = params[4]; // strain 1 half-saturation constant
  double h2 = params[5]; // strain 2 half-saturation constant
  double m0 = params[6]; // baseline mortality rate
  double m1 = params[7]; // drug-induced mortality rate
  double r = params[8]; // drug resistance of strain 1
  double influx = params[9]; // mean resource influx rate
  double outflux = params[10]; // mean resource outflux rate
  double c = params[11]; // colonization/migration rate for bacteria between patches
  int Pt = params[12]; // total number of resource patches
  int P1 = params[13]; // initial number of patches containing strain 1
  int P2 = params[14]; // initial number of patches continaing strain 2
  int N1 = params[15]; // initial population size of strain 1 in occupied patches
  int N2 = params[16]; // initial population size of strain 2 in occupied patches
  double timestep = params[17]; // how frequently to record system information
  double tmax = params[18]; // how long to run simulations
  int global = params[19]; // binary: 0 means migration is local; 1 means migration is global
  double extinctRate = params[20]; // rate that patches go extinct

  // Set up the patches as an NumericMatrix since the total number of patches cannot grow
  // Each patch contains resources and (possibly) bacterial strain 1 and 2
  NumericMatrix Patches(Pt, 3);
  // All patches are assumed to start with resources at carrying capacity (defined by i/o)
  for (int i=0; i < Pt; i++) // set initial resource levels in each patch
    Patches(i,0) = floor(influx/outflux);
  // The first P1 patches (counting up from 0) contain N1 indls of strain 1
  for (int i=0; i < P1; i++) // set initial abundances of strain 1
    Patches(i,1) = N1;
  // Which patches to put strain 2 in depends on whether migration is local or global
  // If global, it doesn't really matter as long as it is likely to avoid strain 1 patches
  // The last P2 patches (counting down from Pt) contain N2 indls of strain 2
  if (global==1) {
    for (int i=(Pt-P2); i < Pt; i++)  // set initial abundances of strain 2
      Patches(i,2) = N2;
  }
  // If local, put them as far apart as possible, so start seeding at patch Pt/2
  // This will generate an error if Pt/2+P2 > Pt
  else {
    for (int i=floor(Pt/2); i < (floor(Pt/2)+P2); i++)
```

```
    Patches(i,2) = N2;
}

// initialize time
double t = 0.0;

// set up a vector containing the times that the metacommunity state should be recorded
NumericVector times(floor(tmax/timestep)+1);
std::iota(times.begin(), times.end(), 0); // creates a vector from 0 to tmax/timestep+1
times = times*timestep; // turns times into a vector from 0 to tmax in units of timestep
int tIter = 0; // what is the current times position?

// initialize storage for the states of the patches at the specified times
List PatchState(times.length());

// get the initial states for all hosts
NumericVector R = Patches(_,0);
NumericVector S1 = Patches(_,1);
NumericVector S2 = Patches(_,2);

// create storage for the events so they aren't created every time through the loop
NumericVector inflow(Pt, influx); // resource inflow rates are always the same
NumericVector outflow, cons, rep1, rep2, death1, death2, move1, move2; // these change with populatio
int nrates = Pt*10; // total number of rate processes equals the number of patches times the number o
NumericVector rates(nrates); // storage for all rates
NumericVector partialRates(nrates); // storage for the fractional rate of each process
double totalRate, rand, totalS1, totalS2;
int event, ind, moveTo;
IntegerVector whichEvent, whichPatch;
// vector for dealing with movement to random patches if movement is global
// patchVector is a vector from 1/Pt to 1 in units of 1/Pt
NumericVector patchVector(Pt);
std::iota(patchVector.begin(), patchVector.end(), 1);
patchVector = patchVector/Pt;

// total population sizes of each strain across all patches
totalS1 = std::accumulate(S1.begin(), S1.end(), 0.0);
totalS2 = std::accumulate(S2.begin(), S2.end(), 0.0);


while(t < tmax && totalS1 > 0.0 && totalS2 > 0.0) {
  // store population information?
  if (t >= times(tIter)) {
    PatchState[tIter] = clone(Patches); // clone() is necessary or every PatchState will become the f
    tIter += 1;
  }

  // compute the rates of all model processes
  outflow = outflux*R; // resource outflow rates in each patch
  cons = 1/Y1*g1/r*R/(h1+R)*S1 + 1/Y2*g2*R/(h2+R)*S2; // resource consumption in each patch
  rep1 = g1/r*R/(h1+R)*S1; // strain 1 replication
  rep2 = g2*R/(h2+R)*S2; // strain 2 replication
  death1 = (m0+m1/(r*r))*S1; // strain 1 death
```

```
death2 = (m0+m1)*S2; // strain 2 death
move1 = c*S1; // strain 1 emigration
move2 = c*S2; // strain 2 emigration

// combine all rates into a single vector
for (int i=0; i < Pt; i++) {
  rates(i) = inflow(i);
  rates(i+Pt) = outflow(i);
  rates(i+2*Pt) = cons(i);
  rates(i+3*Pt) = rep1(i);
  rates(i+4*Pt) = rep2(i);
  rates(i+5*Pt) = death1(i);
  rates(i+6*Pt) = death2(i);
  rates(i+7*Pt) = move1(i);
  rates(i+8*Pt) = move2(i);
  rates(i+9*Pt) = extinctRate;
}
//Rcout << "The rates vector : " << rates << "\n";

// compute the total rate of events across all patches
totalRate = std::accumulate(rates.begin(), rates.end(), 0.0);

// divide each rate by the total rate
partialRates = rates / totalRate;
// cumulative sum the rates to set up the "wheel of fortune"
NumericVector cumsumRates(rates.length());
std::partial_sum(partialRates.begin(), partialRates.end(), cumsumRates.begin());

// generate a random uniform
rand = runif(1)[0];
// identify which event is happening
whichEvent = ifelse(rand > cumsumRates, 1, 0);
event = std::accumulate(whichEvent.begin(), whichEvent.end(), 0);

// increment time
t += rexp(1, totalRate)[0];

// Resource inflow
if (event < Pt) {
  ind = event; // which patch is the event happening in?
  Patches(ind,0) += 1;
}
// Resource outflow
else if (event < (2*Pt)) {
  ind = event-Pt; // which patch is the event happening in?
  Patches(ind,0) -= 1;
}
// Resource consumption
else if (event < (3*Pt)) {
  ind = event-2*Pt; // which patch is the event happening in?
  Patches(ind,0) -= 1;
}
// strain 1 replication
```

```
    else if (event < (4*Pt)) {
      ind = event-3*Pt; // which patch is the event happening in?
      Patches(ind,1) += 1;
    }
    // strain 2 replication
    else if (event < (5*Pt)) {
      ind = event-4*Pt; // which patch is the event happening in?
      Patches(ind,2) += 1;
    }
    // strain 1 death
    else if (event < (6*Pt)) {
      ind = event-5*Pt; // which patch is the event happening in?
      Patches(ind,1) -= 1;
    }
    // strain 2 death
    else if (event < (7*Pt)) {
      ind = event-6*Pt; // which patch is the event happening in?
      Patches(ind,2) -= 1;
    }
    // strain 1 migration
    else if (event < (8*Pt)) {
      ind = event-7*Pt; // which patch is the event happening in?
      Patches(ind,1) -= 1;
      // where does the migrant go?
      // generate a random uniform
      rand = runif(1)[0];
      if (global==1) { // movement is global - emigrant can go to any other patch
        whichPatch = ifelse(rand > patchVector, 1, 0);
        moveTo = std::accumulate(whichPatch.begin(), whichPatch.end(), 0);
        Patches(moveTo,1) += 1;
      } else { // movement is local - emigrant can only go to a neighboring patch (arranged in a ring)
        if (rand < 0.33333333) {
          if (ind > 0)
            moveTo = ind-1;
          else
            moveTo = Pt-1; // if ind is leaving patch 0 for a "smaller" patch, it wraps around to patch
        }
        else if (rand < 0.66666667)
          moveTo = ind; // individual doesn't actually leave
        else {
          if (ind < (Pt-1))
            moveTo = ind+1;
          else
            moveTo = 0; // if ind is leaving patch Pt-1 for a "larger" patch, it wraps around to patch 
        }
        Patches(moveTo,1) += 1;
      }
    }
    // strain 2 migration
    else if (event < (9*Pt)) {
      ind = event-8*Pt; // which patch is the event happening in?
      Patches(ind,2) -= 1;
      // where does the migrant go?
```

```
      // generate a random uniform
      rand = runif(1)[0];
      if (global==1) { // movement is global - emigrant can go to any other patch
        whichPatch = ifelse(rand > patchVector, 1, 0);
        moveTo = std::accumulate(whichPatch.begin(), whichPatch.end(), 0);
        Patches(moveTo,2) += 1;
      } else { // movement is local - emigrant can only go to a neighboring patch (arranged in a ring)
        if (rand < 0.33333333) {
          if (ind > 0)
            moveTo = ind-1;
          else
            moveTo = Pt-1; // if ind is leaving patch 0 for a "smaller" patch, it wraps around to patch
        }
        else if (rand < 0.66666667)
          moveTo = ind; // individual doesn't actually leave
        else {
          if (ind < (Pt-1))
            moveTo = ind+1;
          else
            moveTo = 0; // if ind is leaving patch Pt-1 for a "larger" patch, it wraps around to patch
        }
        Patches(moveTo,2) += 1;
      }
    }
    else { // patch extinction
      ind = event-9*Pt; // which patch is going extinct?
      //Rprintf("%i\n",ind);
      Patches(ind,0) = floor(influx/outflux);
      Patches(ind,1) = 0;
      Patches(ind,2) = 0;
    }

    // get the current states for resources across patches
    R = Patches(_,0);
    S1 = Patches(_,1);
    S2 = Patches(_,2);

    totalS1 = std::accumulate(S1.begin(), S1.end(), 0.0);
    totalS2 = std::accumulate(S2.begin(), S2.end(), 0.0);
  }
  // Get the final system state
  PatchState[tIter] = clone(Patches);
  // return a list
  return PatchState;
}
```

Figure 2:

```
## Stochastic spatial competition model with random patch extinction
sourceCpp("stoch_spatial_comp_3.cpp") ## includes random patch extinction

set.seed(1009409234)
for (iIn in c(50, 100)) {
  for (extinct_rate in c(0.001, 0.005, 0.01)) {
```

```
    ## Vary the growth rate of species 2. When growth_coef = 1, both species have the same growth rate.
    for (growth_coef in 1:20) {
      ## Vary the mortality coefficient from the drugs causing 1x baseline mortality to 20x (set m1=mor
      ## Varying m1 causes the trait, and thus the growth rate, of the resistant strain to change
      for (mort_coef in 16:20) {
        ## Set the deterministic model parameters
        ## Y1,2 = yield
        ## g1,2 = growth rate
        ## h1,2 = half-saturation of Michaelis-Menten function
        ## m0 = baseline mortality rate
        ## m1 = drug-induced mortality rate in the absence of resistance
        ## i = resource input flux
        ## o = resource output rate
        params0 <- c(Y1=0.1, Y2=0.1,
                     g1=5, g2=5*growth_coef,
                     h1=20, h2=20,
                     m0=0.01, m1=0.01*mort_coef,
                     r=0,
                     i=iIn, o=1)
        params0["r"] = sqrt(params0["m1"]/params0["m0"])

        ## Set the stochastic patch extinction model parameters
        ## c = colonization rate
        ## Pt = number of patches
        ## P1,2 = number of patches initially colonized by species 1, 2
        ## timestep = how often to record the system time
        ## tmax = the maximum number of timesteps to allow the simulation to run
        ## global = whether migration is local (0) or global (1)
        ## extinctRate = the rate at which patches stochastically go extinct
        params2 <- c(params0,
                     c(c=0.0001, Pt=20, P1=2, P2=2,
                       N1=10, N2=10,
                       timestep=1, tmax=2000,
                       global=0, ## is migration local or global?
                       extinctRate=extinct_rate)) ## what is the probability a patch goes extinct?

        ## Perform 100 stochastic replicate simulations for each parameter set
        tIn = Sys.time()
        mclapply(1:100,
                 function(x) stoch_spatial_comp_3(params2),
                 mc.cores=10) -> out
        tOut = Sys.time()
        print(paste("resource input =", iIn))
        print(paste("extinction rate =", extinct_rate))
        print(paste("growth coefficient =", growth_coef))
        print(paste("mortality coefficient =", mort_coef))
        print(paste("run time =", tOut-tIn))
        ## Save the resulting data
        saveRDS(out, file=paste0("competition_simulations_growth_coef=",growth_coef,"_mort_coef=",mort_
      }
    }
  }
}
```

```
for (iIn in c(50,100)) {
  for (e_rate in c(0.01, 0.005, 0.001)) {
    print(e_rate)
    results <- expand.grid(rep=1:20, growth_coef=1:20, mort_coef=1:20) %>%
      mutate(.,
             tFinal=0,
             P1=0,
             P2=0,
             meanS1=0,
             meanS2=0)
    for (growth_coef in 1:20) {
      for (mort_coef in 1:20) {
        out <- readRDS(paste0("competition_simulations_growth_coef=",growth_coef,"_mort_coef=",mort_coe
        ## Record the total simulation length, number of patches occupied by each species at the final
        for (rep in 1:100) {
          which.row = which(results$mort_coef==mort_coef &
                              results$growth_coef==growth_coef &
                              results$rep==rep)
          tFinal = sum(unlist(lapply(out[[rep]], function(x) !is.null(x))))
          results$tFinal[which.row] = tFinal
          results$P1[which.row] = sum(out[[rep]][[tFinal]][,2]>0)
          results$P2[which.row] = sum(out[[rep]][[tFinal]][,3]>0)
          results$meanS1[which.row] = mean(out[[rep]][[tFinal]][,2])
          results$meanS2[which.row] = mean(out[[rep]][[tFinal]][,3])
        }
      }
    }
    saveRDS(results, file=paste0("results_competition_fast_vs_resistant_extinct_rate=",e_rate,"_resourc
  }
}

set.seed(1009409234)
for (iIn in c(50,100)) {
  for (extinct_rate in c(0.001, 0.005, 0.01)) {
    ## Vary the yield of species 2. When yield_coef = 1, both species have the same yield. When yield_c
    for (yield_coef in 1:20) {
      ## Vary the mortality coefficient from the drugs causing 1x baseline mortality to 20x
      ## Varying m1 causes the trait, and thus the growth rate, of the resistant strain to change
      for (mort_coef in 1:20) {
        ## Set the deterministic model parameters
        ## Y1,2 = yield
        ## g1,2 = growth rate
        ## h1,2 = half-saturation of Michaelis-Menten function
        ## m0 = baseline mortality rate
        ## m1 = drug-induced mortality rate in the absence of resistance
        ## i = resource input flux
        ## o = resource output rate
        params0 <- c(Y1=0.1, Y2=0.1*yield_coef,
                     g1=5, g2=0.5,
                     h1=20, h2=20,
                     m0=0.01, m1=0.01*mort_coef,
                     r=0,
                     i=iIn, o=1)
```

```
        params0["r"] = sqrt(params0["m1"]/params0["m0"])

        ## Set the stochastic patch extinction model parameters
        ## c = colonization rate
        ## Pt = number of patches
        ## P1,2 = number of patches initially colonized by species 1, 2
        ## timestep = how often to record the system time
        ## tmax = the maximum number of timesteps to allow the simulation to run
        ## global = whether migration is local (0) or global (1)
        ## extinctRate = the rate at which patches stochastically go extinct
        params2 <- c(params0,
                     c(c=0.0001, Pt=20, P1=2, P2=2,
                       N1=10, N2=10,
                       timestep=1, tmax=2000,
                       global=0, ## is migration local or global?
                       extinctRate=extinct_rate)) ## what is the probability a patch goes extinct?

        ## Perform 20 stochastic replicate simulations for each parameter set
        tIn = Sys.time()
        mclapply(1:100,
                 function(x) stoch_spatial_comp_3(params2),
                 mc.cores=10) -> out
        tOut = Sys.time()
        print(paste("resource input =", iIn))
        print(paste("extinction rate =", extinct_rate))
        print(paste("yield coefficient =", yield_coef))
        print(paste("mortality coefficient =", mort_coef))
        print(paste("run time =", tOut-tIn))
        ## Save the resulting data
        saveRDS(out, file=paste0("competition_simulations_yield_coef=",yield_coef,"_mort_coef=",mort_co
      }
    }
  }
}

for (iIn in c(50,100)) {
  for (e_rate in c(0.01, 0.005, 0.001)) {
    print(e_rate)
    results <- expand.grid(rep=1:20, yield_coef=1:20, mort_coef=1:20) %>%
      mutate(.,
             tFinal=0,
             P1=0,
             P2=0,
             meanS1=0,
             meanS2=0)
    ## Vary the yield of species 2. When yield_coef = 1, both species have the same yield. When yield_c
    for (yield_coef in 1:20) {
      ## Vary the mortality coefficient from the drugs causing 1x baseline mortality to 20x
      ## Varying m1 causes the trait, and thus the growth rate, of the resistant strain to change
      for (mort_coef in 1:20) {
        out <- readRDS(file=paste0("competition_simulations_yield_coef=",yield_coef,"_mort_coef=",mort_
        ## Record the total simulation length, number of patches occupied by each species at the final
        for (rep in 1:100) {
```

```
          which.row = which(results$yield_coef==yield_coef &
                             results$mort_coef==mort_coef &
                             results$rep==rep)
          tFinal = sum(unlist(lapply(out[[rep]], function(x) !is.null(x))))
          results$tFinal[which.row] = tFinal
          results$P1[which.row] = sum(out[[rep]][[tFinal]][,2]>0)
          results$P2[which.row] = sum(out[[rep]][[tFinal]][,3]>0)
          results$meanS1[which.row] = mean(out[[rep]][[tFinal]][,2])
          results$meanS2[which.row] = mean(out[[rep]][[tFinal]][,3])
        }
      }
    }
    saveRDS(results, file=paste0("results_competition_efficient_vs_resistant_extinct_rate=",e_rate,"_re
  }
}


##
rbind(
  readRDS("results_competition_fast_vs_resistant_extinct_rate=0.001_resources=100_6-23.RDS") %>%
    mutate(.,
           lh_coef=growth_coef,
           e_rate="Low heterogeneity",
           scenario="Growth advantage",
           prev=meanS2/(meanS1+meanS2)) %>%
    group_by(mort_coef, lh_coef, e_rate, scenario) %>%
    summarize(mean_prev=mean(prev)),
  rbind(
    readRDS("results_competition_fast_vs_resistant_extinct_rate=0.005_resources=100_6-23.RDS") %>%
      mutate(.,
             lh_coef=growth_coef,
             e_rate="Medium heterogeneity",
             scenario="Growth advantage",
             prev=meanS2/(meanS1+meanS2)) %>%
      group_by(mort_coef, lh_coef, e_rate, scenario) %>%
      summarize(mean_prev=mean(prev)),
    readRDS("results_competition_fast_vs_resistant_extinct_rate=0.01_resources=100_6-23.RDS") %>%
      mutate(.,
             lh_coef=growth_coef,
             e_rate="High heterogeneity",
             scenario="Growth advantage",
             prev=meanS2/(meanS1+meanS2)) %>%
      group_by(mort_coef, lh_coef, e_rate, scenario) %>%
      summarize(mean_prev=mean(prev,na.rm=T))
  )
) -> results_s1

rbind(
  readRDS("results_competition_efficient_vs_resistant_extinct_rate=0.001_resources=100_6-23.RDS") %>%
    mutate(.,
           lh_coef=yield_coef,
           e_rate="Low heterogeneity",
           scenario="Efficiency advantage",
```

```
          prev=meanS2/(meanS1+meanS2)) %>%
    group_by(mort_coef, lh_coef, e_rate, scenario) %>%
    summarize(mean_prev=mean(prev)),
  rbind(
    readRDS("results_competition_efficient_vs_resistant_extinct_rate=0.005_resources=100_6-23.RDS") %>%
      mutate(.,
             lh_coef=yield_coef,
             e_rate="Medium heterogeneity",
             scenario="Efficiency advantage",
             prev=meanS2/(meanS1+meanS2)) %>%
      group_by(mort_coef, lh_coef, e_rate, scenario) %>%
      summarize(mean_prev=mean(prev)),
    readRDS("results_competition_efficient_vs_resistant_extinct_rate=0.01_resources=100_6-23.RDS") %>%
      mutate(.,
             lh_coef=yield_coef,
             e_rate="High heterogeneity",
             scenario="Efficiency advantage",
             prev=meanS2/(meanS1+meanS2)) %>%
      group_by(mort_coef, lh_coef, e_rate, scenario) %>%
      summarize(mean_prev=mean(prev))
) -> results_s2

rbind(results_s1, results_s2) -> results
results$e_rate = factor(results$e_rate, levels=c("Low heterogeneity", "Medium heterogeneity", "High hete
results$scenario = factor(results$scenario, levels=c("Growth advantage", "Efficiency advantage"))

ann_df = expand.grid(e_rate = levels(results$e_rate),
                     scenario=levels(results$scenario))
ann_df$label = LETTERS[1:6]
ann_df$x = -Inf
ann_df$y = Inf

png(file="Fig2_competition_predictions.png", height=4, width=5, units='in', res=400)
results %>%
  ggplot(., aes(x=lh_coef, y=mort_coef, z=mean_prev)) +
  geom_contour_filled(aes(fill=after_stat(level_mid))) +
  scale_fill_gradient(low="red",
                      high="blue",
                      limits=c(0,1),
                      oob=scales::squish,
                      guide=guide_colorbar(
                        barwidth=unit(5,"cm"),
                        ticks=TRUE
                      )) +
  geom_text(
    data = ann_df,
    aes(x = x, y = y, label = label),
    hjust = -1,
    vjust = 2,
    color="white",
    inherit.aes = FALSE,
    size = 4
  ) +
```

```
  facet_grid(scenario~e_rate) +
  xlab(expression(S[s]~"relative life history advantage")) +
  ylab(expression("Drug-induced mortality coefficient"~mu)) +
  labs(fill=expression(S[S] ~ "prevalence")) +
  theme_bw() +
  theme(legend.position="bottom",
        legend.key.size=unit(0.3,"cm"),
        legend.text=element_text(size=7),
        legend.title=element_text(size=9),
        legend.spacing.y=unit(1,"mm"),
        legend.margin=margin(1,1,1,1))
dev.off()
```

Figure 3:

```
##
rbind(
  readRDS("results_competition_fast_vs_resistant_extinct_rate=0.001_resources=50_6-23.RDS") %>%
    mutate(.,
           lh_coef=growth_coef,
           e_rate="Low heterogeneity",
           scenario="Growth advantage",
           prev=meanS2/(meanS1+meanS2)) %>%
    group_by(mort_coef, lh_coef, e_rate, scenario) %>%
    summarize(mean_prev=mean(prev)),
  rbind(
    readRDS("results_competition_fast_vs_resistant_extinct_rate=0.005_resources=50_6-23.RDS") %>%
      mutate(.,
             lh_coef=growth_coef,
             e_rate="Medium heterogeneity",
             scenario="Growth advantage",
             prev=meanS2/(meanS1+meanS2)) %>%
      group_by(mort_coef, lh_coef, e_rate, scenario) %>%
      summarize(mean_prev=mean(prev)),
    readRDS("results_competition_fast_vs_resistant_extinct_rate=0.01_resources=50_6-23.RDS") %>%
      mutate(.,
             lh_coef=growth_coef,
             e_rate="High heterogeneity",
             scenario="Growth advantage",
             prev=meanS2/(meanS1+meanS2)) %>%
      group_by(mort_coef, lh_coef, e_rate, scenario) %>%
      summarize(mean_prev=mean(prev,na.rm=T))
  )
) -> results_s1

rbind(
  readRDS("results_competition_efficient_vs_resistant_extinct_rate=0.001_resources=50_6-23.RDS") %>%
    mutate(.,
           lh_coef=yield_coef,
           e_rate="Low heterogeneity",
           scenario="Efficiency advantage",
           prev=meanS2/(meanS1+meanS2)) %>%
    group_by(mort_coef, lh_coef, e_rate, scenario) %>%
    summarize(mean_prev=mean(prev)),
```

```
  rbind(
    readRDS("results_competition_efficient_vs_resistant_extinct_rate=0.005_resources=50_6-23.RDS") %>%
      mutate(.,
             lh_coef=yield_coef,
             e_rate="Medium heterogeneity",
             scenario="Efficiency advantage",
             prev=meanS2/(meanS1+meanS2)) %>%
      group_by(mort_coef, lh_coef, e_rate, scenario) %>%
      summarize(mean_prev=mean(prev)),
    readRDS("results_competition_efficient_vs_resistant_extinct_rate=0.01_resources=50_6-23.RDS") %>%
      mutate(.,
             lh_coef=yield_coef,
             e_rate="High heterogeneity",
             scenario="Efficiency advantage",
             prev=meanS2/(meanS1+meanS2)) %>%
      group_by(mort_coef, lh_coef, e_rate, scenario) %>%
      summarize(mean_prev=mean(prev))
) -> results_s2

rbind(results_s1, results_s2) -> results
results$e_rate = factor(results$e_rate, levels=c("Low heterogeneity", "Medium heterogeneity", "High hete
results$scenario = factor(results$scenario, levels=c("Growth advantage", "Efficiency advantage"))

ann_df = expand.grid(e_rate = levels(results$e_rate),
                     scenario=levels(results$scenario))
ann_df$label = LETTERS[1:6]
ann_df$x = -Inf
ann_df$y = Inf

png(file="FigS3_competition_predictions.png", height=4, width=5, units='in', res=400)
results %>%
  ggplot(., aes(x=lh_coef, y=mort_coef, z=mean_prev)) +
  geom_contour_filled(aes(fill=after_stat(level_mid))) +
  scale_fill_gradient(low="red",
                      high="blue",
                      limits=c(0,1),
                      oob=scales::squish,
                      guide=guide_colorbar(
                        barwidth=unit(5,"cm"),
                        ticks=TRUE
                      )) +
  geom_text(
    data = ann_df,
    aes(x = x, y = y, label = label),
    hjust = -1,
    vjust = 2,
    color="white",
    inherit.aes = FALSE,
    size = 4
  ) +
  facet_grid(scenario~e_rate) +
  xlab(expression(S[s]~"relative life history advantage")) +
  ylab(expression("Drug-induced mortality coefficient"~mu)) +
```

```
  labs(fill=expression(S[S] ~ "prevalence")) +
  theme_bw() +
  theme(legend.position="bottom",
        legend.key.size=unit(0.3,"cm"),
        legend.text=element_text(size=7),
        legend.title=element_text(size=9),
        legend.spacing.y=unit(1,"mm"),
        legend.margin=margin(1,1,1,1))
dev.off()

results = results_s2
results$e_rate = factor(results$e_rate, levels=c("Low heterogeneity", "Medium heterogeneity", "High hete

ann_df = data.frame(e_rate = unique(results$e_rate),
                    label = LETTERS[1:3],
                    x=-Inf,
                    y=Inf)

png(file="Fig3_competition_predictions_low_resources.png", height=3, width=5, units='in', res=400)
ggplot(results, aes(x=lh_coef, y=mort_coef, z=mean_prev)) +
  geom_contour_filled(aes(fill=after_stat(level_mid))) +
  facet_wrap(.~e_rate) +
  scale_fill_gradient(low="red",
                      high="blue",
                      limits=c(0,1),
                      oob=scales::squish,
                      guide=guide_colorbar(
                        barwidth=unit(5,"cm"),
                        ticks=TRUE
                      )) +
  geom_text(
    data = ann_df,
    aes(x = x, y = y, label = label),
    hjust = -1,
    vjust = 2,
    color="white",
    inherit.aes = FALSE,
    size = 4
  ) +
  xlab(expression(S[s]~"relative efficiency advantage")) +
  ylab(expression("Drug-induced mortality"~mu)) +
  labs(fill=expression(S[S] ~ "prevalence")) +
  theme_bw() +
  theme(legend.position="bottom",
        legend.key.size=unit(0.3,"cm"),
        legend.text=element_text(size=7),
        legend.title=element_text(size=9),
        legend.spacing.y=unit(1,"mm"),
        legend.margin=margin(1,1,1,1))
dev.off()
```

C++ code to simulate competition between a strain evolving resistance under a growth–resistance trade-off and a strain with fixed life history:

```cpp
#include <Rcpp.h>
#include <Rmath.h>
#include <numeric>
#include <algorithm>
using namespace Rcpp;

// [[Rcpp::export]]
double lognormal(double mean, double var) {
  double sigma2 = log(var/(mean * mean) + 1.0);
  double sigma = sqrt(sigma2);
  double mu = log(mean) - sigma2/2.0;
  double newInd = R::rlnorm(mu,sigma);
  return newInd;
}

// [[Rcpp::export]]
int eventChooser(NumericVector partialRates, double erand) {
  int event = std::lower_bound(partialRates.begin(), partialRates.end(), erand) - partialRates.begin();
  return event;
}

// [[Rcpp::export]]
double vectorSum(Rcpp::NumericVector x) {
  // std::accumulate takes:
  // 1. An iterator to the beginning of the range (x.begin())
  // 2. An iterator to the end of the range (x.end())
  // 3. An initial value for the sum (0.0 for doubles)
  // 4. An optional binary operation (std::plus<double>() for addition)
  return std::accumulate(x.begin(), x.end(), 0.0, std::plus<double>());
}

// [[Rcpp::export]]
List stoch_spatial_comp_w_evolution(NumericVector params) {

  // Set parameters
  // x1 = parameter for strain evolving under a growth-resistance trade-off
  // x2 = parameter for non-evolving efficient strain
  // Y1, Y2 = yield coefficients
  // g1, g2 = max growth rates
  // h1, h2 = half-saturation constants
  // m0 = baseline mortality rate
  // m1 = drug-induced mortality rate
  // influx = resource influx rate for each patch
  // outflux = resource outflow rate for each patch
  // c = colonization rate
  // Pt = total number of resource patches
  // P1, P2 = number of patches initially colonized by strain 1, strain 2
  // N1, N2 = initial population sizes of strain 1, strain 2 in initially colonized patches
  // timestep = how often to record system state
  // tmax = total time to run simulation
  // global = is colonization local (0) or global (1)
  // extinctRate = stochastic patch extinction rate
  // rvar = additive genetic variance in resistance
```

```
double Y1 = params[0], Y2 = params[1], g1 = params[2], g2 = params[3];
double h1 = params[4], h2 = params[5], m0 = params[6], m1 = params[7];
double influx = params[8], outflux = params[9], c = params[10];
int Pt = params[11], P1 = params[12], P2 = params[13];
int N1 = params[14], N2 = params[15];
double timestep = params[16];
int tmax = params[17];
int global = params[18];
double extinctRate = params[19], rvar = params[20];
double initialResources = floor(influx / outflux);

// Set up a matrix holding the population sizes in each patch
NumericMatrix Patches(Pt, 3);
for (int i = 0; i < Pt; ++i) Patches(i, 0) = initialResources;
for (int i = 0; i < P1; ++i) Patches(i, 1) = N1;
if (global == 1) {
  for (int i = Pt - P2; i < Pt; ++i) Patches(i, 2) = N2;
} else {
  for (int i = Pt / 2; i < Pt / 2 + P2; ++i) Patches(i, 2) = N2;
}

// Set up a list containing the trait values for each individual of strain 1 in each patch
List Traits(Pt);
NumericVector x;
for (int i = 0; i < Pt; i++)
  Traits(i) = x;
for (int i=0; i < P1; i++) {
  // traits are drawn from a lognormal distribution,
  NumericVector traits(N1);
  for (int j=0; j < N1; j++)
    traits[j] = lognormal(1.0, rvar);
  Traits(i) = traits;
}

// create a vector containing the times when the system state should be recorded
double t = 0.0;
int nSteps = floor(tmax / timestep) + 1;
NumericVector times(nSteps);
std::iota(times.begin(), times.end(), 0);
times = times * timestep;
int tIter = 0;

// initialize storage lists for the system state and trait state
List PatchState(nSteps), TraitState(nSteps);

// initialize other storage
NumericVector R = Patches(_, 0), S1 = Patches(_, 1), S2 = Patches(_, 2);
NumericVector inflow(Pt, influx), outflow(Pt), cons(Pt), rep1(Pt), rep2(Pt);
NumericVector death1(Pt), death2(Pt), move1(Pt), move2(Pt);
NumericVector rates(Pt * 10), partialRates(Pt * 10), cumsumRates(Pt * 10);
NumericVector patchVector(Pt);
std::iota(patchVector.begin(), patchVector.end(), 1);
patchVector = patchVector / Pt;
```

```
// compute total S1 and S2
double totalS1 = vectorSum(S1);
double totalS2 = vectorSum(S2);

// simulate the system
while (t < tmax && totalS1 > 0 && totalS2 > 0) {

  // record the state if necessary
  if (t >= times[tIter]) {
    //Rcpp::Rcout << "t = " << t << std::endl;
    PatchState[tIter] = clone(Patches); // use 'clone' to avoid overwriting, since Patches and Traits
    NumericVector meanTraits(Pt);
    for (int i = 0; i < Pt; ++i) {
      NumericVector traits = Traits[i];
      meanTraits[i] = Rcpp::mean(traits);
    }
    TraitState[tIter] = meanTraits;
    //TraitState[tIter] = clone(Traits);
    tIter++;
  }

  // compute mortality rates and replication rates for strain 1, accounting for the trait values
  // for an individual with resistance r, its growth rate is g1/r and its mortality rate is m0+m1/r^2
  // - this trade-off has been proven to produce an ESS under a single-evolution scenario of r = sqrt
  for (int i = 0; i < Pt; ++i) {
    NumericVector traits = Traits[i];
    rep1[i] = 0.0;
    death1[i] = 0.0;
    for (int j = 0; j < traits.size(); ++j) {
      rep1[i] += (g1 / traits[j]) * R[i] / (h1 + R[i]);
      death1[i] += (m0 + m1 / (traits[j] * traits[j]));
    }
  }

  // compute the other rate processes in the model
  outflow = outflux * R; // resource outflow
  rep2 = g2 * R / (h2 + R) * S2; // strain 2 replication
  cons = (1.0 / Y1) * rep1 + (1.0 / Y2) * rep2; // resource consumption
  death2 = (m0 + m1) * S2; // strain 2 mortality
  move1 = c * S1; // strain 1 colonization
  move2 = c * S2; // strain 2 colonization

  // fill in the rates vector with the rates of every process in each patch
  for (int i = 0; i < Pt; ++i) {
    rates[i] = inflow[i];
    rates[i + Pt] = outflow[i];
    rates[i + 2 * Pt] = cons[i];
    rates[i + 3 * Pt] = rep1[i];
    rates[i + 4 * Pt] = rep2[i];
    rates[i + 5 * Pt] = death1[i];
    rates[i + 6 * Pt] = death2[i];
    rates[i + 7 * Pt] = move1[i];
    rates[i + 8 * Pt] = move2[i];
```

```
    rates[i + 9 * Pt] = extinctRate;
  }

  // compute the total event rate
  double totalRate = std::accumulate(rates.begin(), rates.end(), 0.0);
  // stochastic draw for when the next event occurs
  t += rexp(1, totalRate)[0];

  // compute the probability of each event in each patch based on each partial rate and the total rate
  std::transform(rates.begin(), rates.end(), partialRates.begin(), [totalRate](double r) { return r /
  NumericVector cumsumRates(rates.length());
  std::partial_sum(partialRates.begin(), partialRates.end(), cumsumRates.begin());

  //Rcpp::Rcout << "rates = " << rates << std::endl;
  //Rcpp::Rcout << "cumsumRates = " << cumsumRates << std::endl;

  // draw a random number to determine which event occurs
  double erand = R::runif(0.0, 1.0);
  int event = std::lower_bound(cumsumRates.begin(), cumsumRates.end(), erand) - cumsumRates.begin();

  int ind = event % Pt; // determine which patch the event is happening in
  int category = event / Pt; // determine which event is happening

  if (category == 0) Patches(ind, 0) += 1; // resource influx
  else if (category == 1 || category == 2) Patches(ind, 0) -= 1; // resource outflux or consumption
  else if (category == 3) { // strain 1 reproduction
    Patches(ind, 1) += 1;
    NumericVector traits = Traits[ind];
    // choose which individual is replicating based on the individual replication rates
    NumericVector weights = g1 / traits;
    double total = std::accumulate(weights.begin(), weights.end(), 0.0);
    weights = weights / total;
    NumericVector cumsum(weights.size());
    std::partial_sum(weights.begin(), weights.end(), cumsum.begin());
    double r = R::runif(0.0, 1.0);
    int idx = std::lower_bound(cumsum.begin(), cumsum.end(), r) - cumsum.begin();
    // replicate this individual by drawing a random lognormal
    double mean = traits[idx];
    double rNew = lognormal(mean, rvar);
    //Rcpp::Rcout << "old trait = " << mean << std::endl;
    //Rcpp::Rcout << "new trait = " << rNew << std::endl;

    // add the new individual to Traits for this patch
    traits.push_back(rNew);
    Traits[ind] = traits;
  }
  else if (category == 4) Patches(ind, 2) += 1; // strain 2 replication
  else if (category == 5) { // strain 1 mortality
    Patches(ind, 1) -= 1; // remove one individual from the patch
    // choose which individual dies on the basis of its traits
    NumericVector traits = Traits[ind];
    int n = traits.size();
    NumericVector weights(n);
```

```
    for (int ii = 0; ii < n; ++ii) weights[ii] = (m0 + m1 / (traits[ii] * traits[ii]));
    double total = std::accumulate(weights.begin(), weights.end(), 0.0);
    weights = weights / total;
    NumericVector cumsum(weights.size());
    std::partial_sum(weights.begin(), weights.end(), cumsum.begin());
    double r = R::runif(0.0, 1.0);
    int idx = std::lower_bound(cumsum.begin(), cumsum.end(), r) - cumsum.begin();
    // remove this individual from Traits for this patch
    traits.erase(idx);
    Traits[ind] = traits;
  }
  else if (category == 6) Patches(ind, 2) -= 1; // strain 2 mortality
  else if (category == 7 || category == 8) { // migration
    int species = (category == 7) ? 1 : 2; // determine which strain is migrating
    // Rcpp::Rcout << "S1 = " << S1 << std::endl;
    // Rcpp::Rcout << "S2 = " << S2 << std::endl;
    // Rcpp::Rcout << "move1 = " << move1 << std::endl;
    // Rcpp::Rcout << "move2 = " << move2 << std::endl;
    // Rcpp::Rcout << "moving species = " << species << std::endl;
    //
    Patches(ind, species) -= 1; // remove one individual of that strain from the patch
    // identify which patch the individual is moving to
    int moveTo;
    double r = R::runif(0.0, 1.0);
    if (global == 1) {
      moveTo = std::lower_bound(patchVector.begin(), patchVector.end(), r) - patchVector.begin();
    } else {
      if (r < 1.0 / 3.0) moveTo = (ind > 0) ? ind - 1 : Pt - 1;
      else if (r < 2.0 / 3.0) moveTo = ind;
      else moveTo = (ind < Pt - 1) ? ind + 1 : 0;
    }
    // add the individual to the new patch
    Patches(moveTo, species) += 1;
    if (species == 1 && moveTo!=ind) { // if strain 1 moved to a new patch, identify the trait of the
      NumericVector traits = Traits[ind];
      // choose the migrating individual at random
      int idx = R::runif(0.0, traits.size());
      double val = traits[idx];
      // remove it from the current patch
      traits.erase(idx);
      Traits[ind] = traits;
      // and add it to the new patch
      NumericVector newTraits = Traits[moveTo];
      newTraits.push_back(val);
      Traits[moveTo] = newTraits;
    }
  }
  else { // patch extinction
    Patches(ind, 0) = initialResources; // reset the resource and strain densities in this patch
    Patches(ind, 1) = 0;
    Patches(ind, 2) = 0;
    Traits[ind] = NumericVector(); // remove all individuals in the Traits vector from this patch
  }
```

```
    // update the state variables
    R = Patches(_, 0);
    S1 = Patches(_, 1);
    S2 = Patches(_, 2);
    // compute total S1 and S2
    totalS1 = vectorSum(S1);
    totalS2 = vectorSum(S2);

  }

  PatchState[tIter] = clone(Patches);
  NumericVector meanTraits(Pt);
  for (int i = 0; i < Pt; ++i) {
    NumericVector traits = Traits[i];
    meanTraits[i] = Rcpp::mean(traits);
  }
  TraitState[tIter] = meanTraits;
  //TraitState[tIter] = clone(Traits);
  return List::create(Named("PatchState") = PatchState, Named("TraitState") = TraitState);
}
```

Figure 4:

```
sourceCpp("optimized_stoch_spatial_comp_w_evolution.cpp")
set.seed(1009409234)

## For this model, we are competing the evolving fast strain against another fast strain, rather than a
## The point is to provide a contrast to show that (hopefully) competition against an efficient strain i
## much more effective at preventing the evolution of drug resistance than competition against another f
for (e_rate in c(0.01, 0.005, 0.001)) {
  for (mort_coef in seq(1,20,1)) {
    for (growth_coef in seq(1,20,1)) {
      print(paste("e rate", e_rate))
      print(paste("mort", mort_coef))
      print(paste("growth", growth_coef))
      ## influx = 100
      params0 <- c(Y1=0.1, Y2=0.1,
                   g1=5, g2=5*growth_coef,
                   h1=20, h2=20,
                   m0=0.01, m1=0.01*mort_coef,
                   i=100, o=1)
      params <- c(params0,
                  c(c=0.0001, Pt=20, P1=2, P2=2,
                    N1=10, N2=10,
                    timestep=1, tmax=2000,
                    global=0, ## is migration local or global?
                    extinctRate=e_rate,
                    rvar=0.01))
      mclapply(1:20,
               function(i) stoch_spatial_comp_w_evolution(params),
               mc.cores=20) -> output
      saveRDS(output, file=paste0("fast_vs_growth-resistance_trade-off_growth_coef=",growth_coef,"_mort_
    }
  }
```

```
}

for (e_rate in c(0.01, 0.005, 0.001)) {
  results <- expand.grid(rep=1:20, growth_coef=1:20, mort_coef=1:20) %>%
    mutate(.,
           tFinal=0,
           P1=0,
           P2=0,
           meanS1=0,
           meanS2=0)

  ## Vary the mortality coefficient from the drugs causing 1x baseline mortality to 20x
  ## Varying m1 causes the trait, and thus the growth rate, of the resistant strain to change
  for (growth_coef in 1:20) {
    for (mort_coef in 1:20) {
      out <- readRDS(paste0("fast_vs_growth-resistance_trade-off_growth_coef=",growth_coef,"_mort_coef=
      ## Record the total simulation length, number of patches occupied by each species at the final ti
      for (rep in 1:20) {
        which.row = which(results$mort_coef==mort_coef &
                            results$growth_coef==growth_coef &
                            results$rep==rep)
        tFinal = sum(unlist(lapply(out[[rep]][[1]], function(x) !is.null(x))))
        results$tFinal[which.row] = tFinal
        results$P1[which.row] = sum(out[[rep]][[1]][[tFinal]][,2]>0)
        results$P2[which.row] = sum(out[[rep]][[1]][[tFinal]][,3]>0)
        results$meanS1[which.row] = mean(out[[rep]][[1]][[tFinal]][,2])
        results$meanS2[which.row] = mean(out[[rep]][[1]][[tFinal]][,3])
      }
    }
  }
  saveRDS(results, file=paste0("results_evolution_fast_vs_resistant_extinct_rate=",e_rate,"_12-5.RDS"))
}

rbind(
  readRDS("results_evolution_fast_vs_resistant_extinct_rate=0.001_12-5.RDS") %>%
    mutate(.,
           lh_coef=growth_coef,
           e_rate="Low heterogeneity",
           scenario="Growth advantage",
           prev=meanS2/(meanS1+meanS2)) %>%
    group_by(mort_coef, lh_coef, e_rate, scenario) %>%
    summarize(mean_prev=mean(prev,na.rm=T)),
  rbind(
    readRDS("results_evolution_fast_vs_resistant_extinct_rate=0.005_12-5.RDS") %>%
      mutate(.,
             lh_coef=growth_coef,
             e_rate="Medium heterogeneity",
             scenario="Growth advantage",
             prev=meanS2/(meanS1+meanS2)) %>%
      group_by(mort_coef, lh_coef, e_rate, scenario) %>%
      summarize(mean_prev=mean(prev,na.rm=T)),
    readRDS("results_evolution_fast_vs_resistant_extinct_rate=0.01_12-5.RDS") %>%
      mutate(.,
```

```
            lh_coef=growth_coef,
            e_rate="High heterogeneity",
            scenario="Growth advantage",
            prev=meanS2/(meanS1+meanS2)) %>%
      group_by(mort_coef, lh_coef, e_rate, scenario) %>%
      summarize(mean_prev=mean(prev,na.rm=T))
  )
) -> results_s1

for (e_rate in c(0.01, 0.005, 0.001)) {
  for (mort_coef in seq(1,20,1)) {
    for (yield_coef in seq(1,20,1)) {
      print(paste("e rate", e_rate))
      print(paste("mort", mort_coef))
      print(paste("yield", yield_coef))
      ## influx = 100
      params0 <- c(Y1=0.1, Y2=0.1*yield_coef,
                   g1=5, g2=0.5,
                   h1=20, h2=20,
                   m0=0.01, m1=0.01*mort_coef,
                   i=100, o=1)
      params <- c(params0,
                  c(c=0.0001, Pt=20, P1=2, P2=2,
                    N1=10, N2=10,
                    timestep=1, tmax=2000,
                    global=0, ## is migration local or global?
                    extinctRate=e_rate,
                    rvar=0.01))
      mclapply(1:20,
               function(i) stoch_spatial_comp_w_evolution(params),
               mc.cores=10) -> output
      saveRDS(output, file=paste0("efficient_vs_growth-resistance_trade-off_yield_coef=",yield_coef,"_mo
    }
  }
}

for (e_rate in c(0.005, 0.01)) {
  print(e_rate)
  results <- expand.grid(rep=1:20, yield_coef=1:20, mort_coef=1:20) %>%
    mutate(.,
           tFinal=0,
           P1=0,
           P2=0,
           meanS1=0,
           meanS2=0)
  for (yield_coef in 1:20) {
    ## Vary the mortality coefficient from the drugs causing 1x baseline mortality to 20x
    ## Varying m1 causes the trait, and thus the growth rate, of the resistant strain to change
    for (mort_coef in 1:20) {
      out <- readRDS(file=paste0("efficient_vs_growth-resistance_trade-off_yield_coef=",yield_coef,"_mo
      ## Record the total simulation length, number of patches occupied by each species at the final ti
      for (rep in 1:20) {
        which.row = which(results$yield_coef==yield_coef &
```

```
                              results$mort_coef==mort_coef &
                              results$rep==rep)
        tFinal = sum(unlist(lapply(out[[rep]][[2]], function(x) !is.null(x))))
        results$tFinal[which.row] = tFinal
        results$P1[which.row] = sum(out[[rep]][[1]][[tFinal]][,2]>0)
        results$P2[which.row] = sum(out[[rep]][[1]][[tFinal]][,3]>0)
        results$meanS1[which.row] = mean(out[[rep]][[1]][[tFinal]][,2])
        results$meanS2[which.row] = mean(out[[rep]][[1]][[tFinal]][,3])
      }
    }
  }
  saveRDS(results, file=paste0("results_efficient_vs_resistant_extinct_rate=",e_rate,"_9-8.RDS"))
}
e_rate = 0.001
results <- expand.grid(rep=1:20, yield_coef=1:20, mort_coef=1:5) %>%
  mutate(.,
         tFinal=0,
         P1=0,
         P2=0,
         meanS1=0,
         meanS2=0)
for (yield_coef in 1:20) {
  ## Vary the mortality coefficient from the drugs causing 1x baseline mortality to 20x
  ## Varying m1 causes the trait, and thus the growth rate, of the resistant strain to change
  for (mort_coef in 1:5) {
    out <- readRDS(file=paste0("efficient_vs_growth-resistance_trade-off_yield_coef=",yield_coef,"_mort_
    ## Record the total simulation length, number of patches occupied by each species at the final time
    for (rep in 1:20) {
      which.row = which(results$yield_coef==yield_coef &
                        results$mort_coef==mort_coef &
                        results$rep==rep)
      tFinal = sum(unlist(lapply(out[[rep]][[2]], function(x) !is.null(x))))
      results$tFinal[which.row] = tFinal
      results$P1[which.row] = sum(out[[rep]][[1]][[tFinal]][,2]>0)
      results$P2[which.row] = sum(out[[rep]][[1]][[tFinal]][,3]>0)
      results$meanS1[which.row] = mean(out[[rep]][[1]][[tFinal]][,2])
      results$meanS2[which.row] = mean(out[[rep]][[1]][[tFinal]][,3])
    }
  }
}
saveRDS(results, file=paste0("results_efficient_vs_resistant_extinct_rate=",e_rate,"_9-8.RDS"))

rbind(
  rbind(
    readRDS("results_efficient_vs_resistant_extinct_rate=0.001_9-8.RDS") %>%
      mutate(.,
             lh_coef=yield_coef,
             e_rate="Low heterogeneity",
             scenario="Efficiency advantage",
             prev=meanS2/(meanS1+meanS2)) %>%
      group_by(mort_coef, lh_coef, e_rate, scenario) %>%
      summarize(mean_prev=mean(prev)),
    expand.grid(mort_coef=6:20,
```

```
                lh_coef=1:20,
                e_rate="Low heterogeneity",
                scenario="Efficiency advantage",
                mean_prev=0)),
  rbind(
    readRDS("results_efficient_vs_resistant_extinct_rate=0.005_9-8.RDS") %>%
      mutate(.,
             lh_coef=yield_coef,
             e_rate="Medium heterogeneity",
             scenario="Efficiency advantage",
             prev=meanS2/(meanS1+meanS2)) %>%
      group_by(mort_coef, lh_coef, e_rate, scenario) %>%
      summarize(mean_prev=mean(prev)),
    readRDS("results_efficient_vs_resistant_extinct_rate=0.01_9-8.RDS") %>%
      mutate(.,
             lh_coef=yield_coef,
             e_rate="High heterogeneity",
             scenario="Efficiency advantage",
             prev=meanS2/(meanS1+meanS2)) %>%
      group_by(mort_coef, lh_coef, e_rate, scenario) %>%
      summarize(mean_prev=mean(prev)))
) -> results_s2

rbind(results_s1, results_s2) -> results
results$e_rate = factor(results$e_rate, levels=c("Low heterogeneity", "Medium heterogeneity", "High hete
results$scenario = factor(results$scenario, levels=c("Growth advantage", "Efficiency advantage"))

ann_df = expand.grid(e_rate = levels(results$e_rate),
                     scenario=levels(results$scenario))
ann_df$label = LETTERS[1:6]
ann_df$x = -Inf
ann_df$y = Inf

png(file="Fig4_evolution_predictions.png", height=4, width=5, units='in', res=400)
results %>%
  ggplot(., aes(x=lh_coef, y=mort_coef, z=mean_prev)) +
  geom_contour_filled(aes(fill=after_stat(level_mid))) +
  scale_fill_gradient(low="red",
                      high="blue",
                      limits=c(0,1),
                      oob=scales::squish,
                      guide=guide_colorbar(
                        barwidth=unit(5,"cm"),
                        ticks=TRUE
                      )) +
  geom_text(
    data = ann_df,
    aes(x = x, y = y, label = label),
    hjust = -1,
    vjust = 2,
    color="white",
    inherit.aes = FALSE,
    size = 4
```

```
  ) +
  facet_grid(scenario~e_rate) +
  xlab(expression(S[s]~"relative life history advantage")) +
  ylab(expression("Drug-induced mortality coefficient"~mu)) +
  labs(fill=expression(S[S] ~ "prevalence")) +
  theme_bw() +
  theme(legend.position="bottom",
        legend.key.size=unit(0.3,"cm"),
        legend.text=element_text(size=7),
        legend.title=element_text(size=9),
        legend.spacing.y=unit(1,"mm"),
        legend.margin=margin(1,1,1,1))
dev.off()
```

Figure 5:

```
sourceCpp("optimized_stoch_spatial_comp_w_evolution.cpp")

for (e_rate in c(0.01, 0.005, 0.001)) {
  for (mort_coef in seq(1,20,1)) {
    for (yield_coef in seq(1,20,1)) {
      print(paste("e rate", e_rate))
      print(paste("mort", mort_coef))
      print(paste("yield", yield_coef))
      ## influx = 50
      params0 <- c(Y1=0.1, Y2=0.1*yield_coef,
                   g1=5, g2=0.5,
                   h1=20, h2=20,
                   m0=0.01, m1=0.01*mort_coef,
                   i=50, o=1)
      params <- c(params0,
                  c(c=0.0001, Pt=20, P1=2, P2=2,
                    N1=10, N2=10,
                    timestep=1, tmax=2000,
                    global=0, ## is migration local or global?
                    extinctRate=e_rate,
                    rvar=0.01))
      mclapply(1:20,
               function(i) stoch_spatial_comp_w_evolution(params),
               mc.cores=10) -> output
      saveRDS(output, file=paste0("efficient_vs_growth-resistance_trade-off_yield_coef=",yield_coef,"_m
    }
  }
}

for (e_rate in c(0.001, 0.005, 0.01)) {
  print(e_rate)
  results <- expand.grid(rep=1:20, yield_coef=1:20, mort_coef=1:20) %>%
    mutate(.,
           tFinal=0,
           P1=0,
           P2=0,
           meanS1=0,
           meanS2=0)
```

```
  for (yield_coef in 1:20) {
    ## Vary the mortality coefficient from the drugs causing 1x baseline mortality to 20x
    ## Varying m1 causes the trait, and thus the growth rate, of the resistant strain to change
    for (mort_coef in 1:20) {
      out <- readRDS(file=paste0("efficient_vs_growth-resistance_trade-off_yield_coef=",yield_coef,"_mor
      ## Record the total simulation length, number of patches occupied by each species at the final ti
      for (rep in 1:20) {
        which.row = which(results$yield_coef==yield_coef &
                            results$mort_coef==mort_coef &
                            results$rep==rep)
        tFinal = sum(unlist(lapply(out[[rep]][[2]], function(x) !is.null(x))))
        results$tFinal[which.row] = tFinal
        results$P1[which.row] = sum(out[[rep]][[1]][[tFinal]][,2]>0)
        results$P2[which.row] = sum(out[[rep]][[1]][[tFinal]][,3]>0)
        results$meanS1[which.row] = mean(out[[rep]][[1]][[tFinal]][,2])
        results$meanS2[which.row] = mean(out[[rep]][[1]][[tFinal]][,3])
      }
    }
  }
  saveRDS(results, file=paste0("results_efficient_vs_resistant_extinct_rate=",e_rate,"_8-29.RDS"))
}

rbind(
  readRDS("results_efficient_vs_resistant_extinct_rate=0.001_8-29.RDS") %>%
    mutate(.,
           lh_coef=yield_coef,
           e_rate="Low heterogeneity",
           scenario="Efficiency advantage",
           prev=meanS2/(meanS1+meanS2)) %>%
    group_by(mort_coef, lh_coef, e_rate, scenario) %>%
    summarize(mean_prev=mean(prev)),
  rbind(
    readRDS("results_efficient_vs_resistant_extinct_rate=0.005_8-29.RDS") %>%
      mutate(.,
             lh_coef=yield_coef,
             e_rate="Medium heterogeneity",
             scenario="Efficiency advantage",
             prev=meanS2/(meanS1+meanS2)) %>%
      group_by(mort_coef, lh_coef, e_rate, scenario) %>%
      summarize(mean_prev=mean(prev)),
    readRDS("results_efficient_vs_resistant_extinct_rate=0.01_8-29.RDS") %>%
      mutate(.,
             lh_coef=yield_coef,
             e_rate="High heterogeneity",
             scenario="Efficiency advantage",
             prev=meanS2/(meanS1+meanS2)) %>%
      group_by(mort_coef, lh_coef, e_rate, scenario) %>%
      summarize(mean_prev=mean(prev)))
) -> results

results$e_rate = factor(results$e_rate, levels=c("Low heterogeneity", "Medium heterogeneity", "High hete

ann_df = data.frame(e_rate = unique(results$e_rate),
```

```
                    label = LETTERS[1:3],
                    x=-Inf,
                    y=Inf)

png(file="Fig5_evolution_predictions_low_resources.png", height=3, width=5, units='in', res=400)
ggplot(results, aes(x=lh_coef, y=mort_coef, z=mean_prev)) +
  geom_contour_filled(aes(fill=after_stat(level_mid))) +
  facet_wrap(.~e_rate) +
  scale_fill_gradient(low="red",
                      high="blue",
                      limits=c(0,1),
                      oob=scales::squish,
                      guide=guide_colorbar(
                        barwidth=unit(5,"cm"),
                        ticks=TRUE
                      )) +
  geom_text(
    data = ann_df,
    aes(x = x, y = y, label = label),
    hjust = -1,
    vjust = 2,
    color="white",
    inherit.aes = FALSE,
    size = 4
  ) +
  xlab(expression(S[s]~"relative efficiency advantage")) +
  ylab(expression("Drug-induced mortality"~mu)) +
  labs(fill=expression(S[S] ~ "prevalence")) +
  theme_bw() +
  theme(legend.position="bottom",
        legend.key.size=unit(0.3,"cm"),
        legend.text=element_text(size=7),
        legend.title=element_text(size=9),
        legend.spacing.y=unit(1,"mm"),
        legend.margin=margin(1,1,1,1))
dev.off()
```
